# Germline and somatic genetic variants in the p53 pathway interact to affect cancer risk, progression and drug response

**DOI:** 10.1101/835918

**Authors:** Ping Zhang, Isaac Kitchen-Smith, Lingyun Xiong, Giovanni Stracquadanio, Katherine Brown, Philipp Richter, Marsha Wallace, Elisabeth Bond, Natasha Sahgal, Samantha Moore, Svanhild Nornes, Sarah De Val, Mirvat Surakhy, David Sims, Xuting Wang, Douglas A. Bell, Jorge Zeron-Medina, Yanyan Jiang, Anderson Ryan, Joanna Selfe, Janet Shipley, Siddhartha Kar, Paul Pharoah, Chey Loveday, Rick Jansen, Lukasz F. Grochola, Claire Palles, Andrew Protheroe, Val Millar, Daniel Ebner, Meghana Pagadala, Sarah P. Blagden, Tim Maughan, Enric Domingo, Ian Tomlinson, Clare Turnbull, Hannah Carter, Gareth Bond

## Abstract

Insights into oncogenesis derived from cancer susceptibility loci could facilitate better cancer management and treatment through precision oncology. However, therapeutic applications have thus far been limited by our current lack of understanding regarding both their interactions with somatic cancer driver mutations and their influence on tumorigenesis. Here, by integrating germline datasets relating to cancer susceptibility with tumour data capturing somatically-acquired genetic variation, we provide evidence that single nucleotide polymorphism (SNPs) and somatic mutations in the p53 tumor suppressor pathway can interact to influence cancer development, progression and treatment response. We go on to provide human genetic evidence of a tumor-promoting role for the pro-survival activities of p53, which supports the development of more effective therapy combinations through their inhibition in cancers retaining wild-type p53.

**Significance:** We describe significant interactions between heritable and somatic genetic variants in the p53 pathway that affect cancer susceptibility, progression and treatment response. Our results offer evidence of how cancer susceptibility SNPs can interact with cancer driver genes to affect cancer progression and identify novel therapeutic targets.

## Introduction

Efforts to characterize the somatic alterations that drive oncogenesis have led to the development of targeted therapies, facilitating precision approaches that condition treatment on knowledge of the tumor genome, and improving outcomes for many cancer patients (1,2). However, such targeted therapies are associated with variable responses, eventual high failure rates and the development of drug resistance. Somatic genetic heterogeneity among tumors is a major factor contributing to differences in disease progression and therapeutic response (1). The maps of common germline genetic variants that associate with disease susceptibility allow us to generate and test biological hypotheses, characterize regulatory mechanisms by which variants contribute to disease, with the aim of integrating the results into the clinic. However, there are challenges in harnessing of susceptibility loci for target identification for cancer, including limitations in (i) exposition of causative variants within susceptibility loci, (ii) understanding of interactions of susceptibility variants with somatic driver mutations, and (iii) mechanistic insights into their influence on cellular behaviors during and after the evolution of somatic cancer genomes (3–5).

A key cancer signaling pathway known to harbor multiple germline and somatic variants associated with cancer susceptibility is the p53 tumor suppressor pathway (6). It is a stress response pathway that maintains genomic integrity and is among the most commonly perturbed pathways in cancer, with somatic driver mutations found in the *TP53* gene in more than 50% of cancer genomes (7). Loss of the pathway and/or the gain of pro-cancer mutations can lead to cellular transformation and tumorigenesis (8). Once cancer has developed, the p53 pathway is important in mediating cancer progression and the response to therapy, as its anti-cancer activities can be activated by many genotoxic anticancer drugs (9). These drugs are more effective in killing cancers with wild-type p53 relative to mutant p53 (10,11). While both germline and somatic alterations to the p53 pathway are known to promote tumorigenesis, the extent to which such variants cooperate to alter pathway activity and the effects on response to therapy remain poorly understood.

In general, p53 mutations drive cancer through loss of wild-type function, dominant negative and gain-of-function activities which have been demonstrated to confer pro-cancer activities such as metastasis, altered energy metabolism, and replicative immortality (12–14). Mutations are primarily missense mutations that affect p53’s ability to bind to DNA in a sequence-specific manner and regulate transcription of its target genes. Some of these same *TP53* mutations when found constitutionally result in Li-Fraumeni Syndrome: a syndrome comprising dramatic increase in cancer risk in many tissues types. Although targeting driver mutations in tumor suppressors has been challenging, the high abundance of p53 mutations in cancer has motivated the development of small molecules that aim to reactivate mutant p53 to increase sensitivities to DNA-damaging therapies or inhibit gain-of function activities (15).

Somatic driver mutations in other p53 pathway genes are also current drug targets. In a sub-set of p53 wild-type cancers, p53 signaling can be attenuated through somatic driver events that alter key p53 regulators. For example, the MDM2 oncogene is amplified in a variety of cancers. Its amplification results in decreased p53-mediated tumor suppression, increased cancer susceptibility, and the reduction of selection pressures for somatic p53 mutations (16). Moreover, cancer cells with amplified MDM2 and wild-type p53 have an attenuated p53-mediated DNA damage response (17). Thus, amplification of MDM2 is a promising target for treatment, in combination with DNA-damaging therapies (15,18).

Most studies have separately examined the consequences of somatic and germline variation affecting p53 activity to understand their roles in disease risk, progression or response to therapy. Here we hypothesize that cancer-associated germline variants (single nucleotide polymorphisms, SNPs) interact with p53 somatic driver mutations to modify cancer risk, progression and potential to respond to therapy. With a focus on cancer-associated SNPs with the potential to directly influence p53 activity, we provide supportive evidence for this hypothesis, and go on to demonstrate their ability to discover candidate drug targets.

## Results

### 1. p53 regulatory cancer risk SNPs associate with subtype heterogeneity

We first explored whether cancer susceptibility SNPs could influence the frequency of somatic mutation of *TP53* of tumors arising in carriers. It is known that key regulatory pathway genes and stress signals, which can regulate wild-type p53 levels and tumor suppressive activities, can also regulate mutant p53, including its oncogenic activities (19–21). Thus, we reasoned that key p53 regulatory genes could have SNPs that modify the ability of mutant p53 to drive cancer and of wild type (WT) p53 to suppress it. If true, these SNPs could associate with allelic-differences in susceptibility to both WT and mutant p53 cancers, but the direction of their associations with risk would be different (heterogeneity risk SNPs) (**Fig. 1A**).

**Figure 1.**
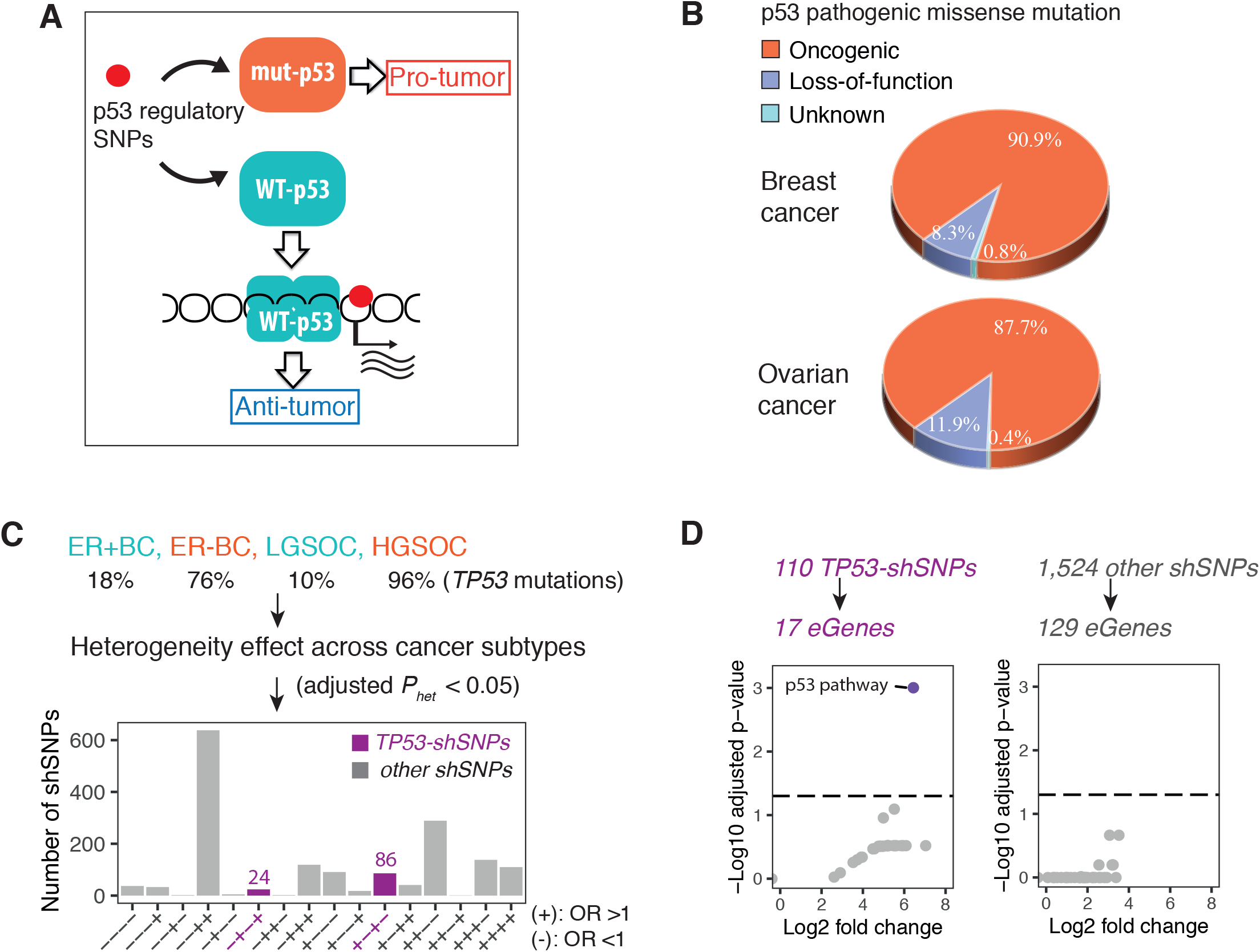
p53 regulatory cancer risk SNPs associate with subtype heterogeneity risk. (A) A proposed model of how p53 regulatory SNPs could modify the ability of mutant p53 to drive cancer and of wild type (WT) p53 to suppress it. (B) Pie charts of the percentages of oncogenic and loss-of-function p53 mutations found amongst all known pathogenic p53 missense mutations in breast and ovarian cancers. (C) A bar plot of the number of SNPs associated with subtype heterogeneity (adjusted *P*_*het*_ < 0.05) across breast and ovarian cancer subtypes (shSNPs). The shSNPs are binned in groups based on the allelic differences in risk found in the various subtypes. For example, +−+−indicates an allele of the SNP associated with an increased risk for ER+BC and LGSOC (OR>1), but lower risk in ER-BC and HGSOC (OR<1). Those shSNPs with allelic differences in risk in the various subtypes that is consistent with p53 mutation frequencies are highlighted in purpled and labeled p53-shSNPs. (D) A scatter plot of the fold enrichment of subtype heterogeneity eGenes amongst all KEGG annotated signaling pathways relative to all eGenes in the genome. The x-axis in a log2 scale, and the adjusted p-value on the y-axis is a −log10 scale. The p53 pathway is in purple and the other 185 annotated KEGG pathways are in grey. The horizontal dashed lines represent the FDR-adjusted p value of 0.05.

To test this, we first sought to identify cancer risk SNPs that are potential modifiers of p53 activity from existing GWAS and eQTL data. Specifically, we identified cancer risk-associated SNPs determined by GWAS, which have also been found to associate with differential expression levels of p53 pathway genes in eQTL databases. There are currently 1,225 cancer GWAS lead SNPs (p < 5e-08) in the GWAS database, which are in linkage disequilibrium (LD) with 27,367 proxies (r2 > 0.8 in EUR). In the three largest expression quantitative trait loci (cis-eQTLs) datasets, 15,406 of these cancer risk SNPs (lead SNPs and proxies) reside in eQTLs (eSNPs) associating with allelic differences in expression levels of 1,438 genes (eGenes) in at least one tissue/cell type (22–24) (**Supplementary Table S1**). When the eGenes are attributed to well-annotated cellular pathways (KEGG pathway database, Methods), we find that p53 pathway genes are over-represented relative to all other annotated pathways in the database (p= 5.5−06, adjust p = 7.8e-05; **Supplementary Fig. 1A**), similar to the results found in a previously published study (6). The p53 pathway eGenes include the *TP53* gene itself, as well as key regulator genes (*MDM4*; *ATM*; *CHEK2*; *CDKN2A*) and key effector genes (*CASP8*; *CDKN1A*; *FAS*; *PIDD*; *CCNE1*; *CCND1*; *SESN1*; *PMAIP1*).

Next, we sought to identify a population of heterogeneity risk SNPs in cancers that associate with disease subtypes that differ substantially in p53 mutation frequencies and for which susceptibility GWAS data was available. 18% of estrogen receptor positive breast cancers (ER+BC) mutate p53, in contrast to 76% of estrogen receptor negative breast cancers (ER−BC) (25). Similarly, less than 10% of low-grade serous ovarian cancers (LGSOC) mutate p53, in contrast to 96% of high grade serous ovarian cancers (HGSOC) (26). Over 85% of p53 pathogenic missense mutations in breast and ovarian cancers are oncogenic (either dominant negative or gain-of-function) (**Fig. 1B**) (see Methods). We analyzed data from 90,969 breast cancer patients of European ancestry (69,501 ER-pos BC, 21,468 ER-neg BC) (27) and 105,974 controls, and 14,049 ovarian cancer patients of European ancestry (1,012 LGSOC, 13,037 HGSOC) and 40,941 controls (28). We found that, of the 15,406 cancer risk eSNPs, 1,634 showed significant subtype heterogeneity after correction for multiple hypothesis testing (Bonferonni adjusted *P*_*het*_ <0.05; **Supplementary Table S2**) across the four subtypes (ER+BC, ER−BC, LGSOC, HGSOC) (subtype heterogeneity SNPs, shSNPs). (**Fig. 1C**). For 110 out of the 1,634 shSNPs, the directions of the allelic-associations with risk were consistent with the p53 mutational frequencies of the breast and ovarian subtypes (*TP53*-relevant subtype heterogeneity SNPs, *TP53*-shSNPs; **Fig. 1C**, purple bars). That is, the alleles of these SNPs that are associated with increased cancer risk (OR>1) in the subtypes with low p53 mutation frequencies (ER+BC and LGSOC), are associated with decreased cancer risk (OR<1) in the subtypes with high p53 mutation frequencies (ER−BC and HGSOC), and vice versa.

The 110 *TP53*-shSNPs are eSNPs for 17 eGenes and the remaining 1,524 shSNPs (other-shSNPs) are eSNPs for 129 eGenes. We reasoned that if key p53 regulatory genes have SNPs that modify the ability of mutant p53 to drive cancer and of wild type (WT) p53 to suppress it, p53 pathway genes could be enriched amongst the 17 eGenes defined by the TP53-shSNPs. Indeed, the 17 eGenes are only significantly enriched in p53 pathway genes and no other annotated pathway (KEGG: 87.0-fold, adjusted p = 9.9e-04; **Fig. 1D**, left panel). Importantly, no such enrichment of p53 pathway genes, or any other pathways, is seen in the 129 eGenes defined by the other-shSNPs (Fig. 1D, right panel). The p53 pathway eGenes include the *TP53* gene itself, as well as key p53 regulator genes (*MDM4* and *ATM*). Thus, key p53 pathway genes harbor cancer-associated regulatory SNPs, which significantly associate with subtype heterogeneity in a manner that follows p53 mutational frequencies: 44 eSNPs in *ATM*, 33 eSNPs in *MDM4* and 3 eSNPs in *TP53.* All SNPs in each gene are in LD (r^2^ and/or d’ >0.9 in Europeans) (**Fig. 2A**; **Supplementary Table S3**).

**Figure 2.**
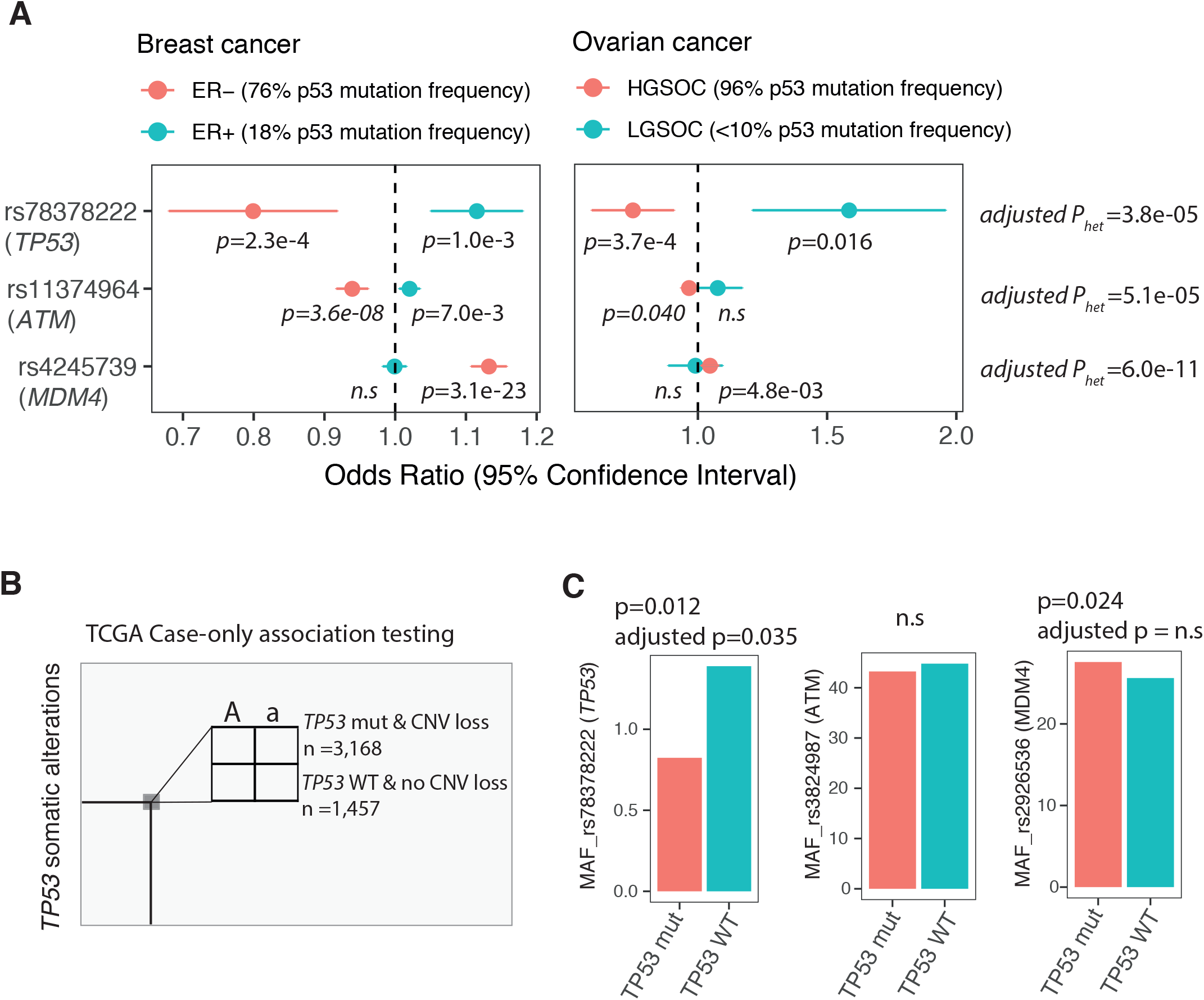
p53 regulatory cancer risk SNPs associate with somatic *TP53* mutational status. (A) Forest plots illustrating the associations of the three p53 regulatory SNPs with breast cancer (left) and ovarian cancer (right) subtype heterogeneity. The odd ratios (OR) are plotted for each SNP and subtype and the error bars represent the associated 95% confidence intervals (CI). (B) A schematic overview of the association testing between the three p53 regulatory risk SNPs and p53 mutational status in 4,625 tumors (TCGA). (C) Bar plots of the minor allele frequencies (MAFs) of the three p53 regulatory SNPs in patients with either WT TP53 tumors or mut TP53 tumors.

### 2. p53 regulatory cancer risk SNPs associate with somatic *TP53* mutational status

Each of these three loci have been previously found to associate with differential risk for at least one cancer in the broader population (29–32), and the above analysis provides evidence that they significantly associate with subtype heterogeneity in a manner that follows p53 mutational frequencies. This observation supports a persistent effect for p53 pathway cancer risk SNPs on tumors through a possible influence on whether or not a tumour contains a somatically mutated *TP53* locus. In order to seek further and more direct support of this possibility, we performed similar analyses of these three loci in a cohort of 7,021 patients of European origin diagnosed with 31 different cancers and for whom the p53 mutational status of their cancers could be determined (The Cancer Genome Atlas, TCGA).

In this cohort, 35.8% of patients have at least one pathogenic p53 mutation in their cancers, 37.8% have p53 copy-number loss (CNV loss), and 20.8% have both (**Supplementary Table S4**). We partitioned the patients into two groups based on the presence or absence of the p53 somatic alteration (mutation and CNV loss versus WT and no CNV loss). We hypothesized that if an allele of a given SNP was found to associate with increased risk in cancer subtypes with lower p53 frequencies, it will be more frequent in those patients with wild type TP53 cancers/tumours, and vice versa. Thus, we performed association testing between the three loci and p53 somatic alterations using a Fisher exact test (**Fig. 2B**, one-sided). For each locus, we performed association testing using the SNP that showed the strongest associations with subtype heterogeneity and, for which, genotype information was available. Interestingly, two of these three SNPs associated with allelic differences in minor allele frequencies between the groups of patients with either p53 WT or mutant tumours (*TP53* SNP and the *MDM4* SNP; **Fig. 2C**). Importantly, the association of the *TP53* SNP, rs78378222, remains significant even after correction for multiple hypothesis testing (Bonferonni adjusted p = 0.035; **Fig. 2C**). For this SNP, the minor C-allele is associated with increased cancer risk in ER+BC and LGSOC (less p53 mutations), but decreased cancer risk in ER−BC and HGSOC (more p53 mutations) (**Fig. 2A**). This is in line with the associations found with p53 mutational status, whereby the C-allele is more frequent in TP53 WT tumors (**Fig. 2C**). Together, these observations lend further support to a persistent effect for p53 pathway cancer risk-associated SNPs on tumors through a possible influence on whether or not a tumor contains a somatically mutated TP53 locus.

### 3. A p53 regulatory cancer risk SNP can affect wild type and mutant p53 in tumors

The TP53 SNP, rs78378222, resides in the 3’-UTR (p53 poly(A) SNP). The minor C-allele has been previously found to associate with lower p53 mRNA levels in blood samples (33). Indeed, when we examine all cellular transcripts using genotype and gene expression data from 4,896 peripheral blood samples, the p53 poly(A) SNP only associates with allelic differences in p53 RNA levels and no other transcripts, whereby the minor C-allele associates with less p53 transcripts (p=2.0e-25, beta=−0.62; **Fig. 3A**). To investigate the activity of this SNP in tumors, we analyzed expression data from 3,248 tumors from the TCGA cohort, for which both germline and somatic genetic data was available and no somatic copy number variation of p53 could be detected. Similar to results obtained in the blood samples, we observed a significant association of the minor C-allele with lower p53 expression levels in the tumors (p=1.7e-04, beta=−0.37; **Fig. 3B**). To test if the C-allele associates with lower levels of both wild type and mutant p53, we divided the tumors into three groups based on their respective somatic p53 mutational status (**Fig. 3C** and **Supplementary Table** S4). We found 2,521 tumors with wild type p53 genes, 448 with missense mutations, and, of those, 389 with oncogenic missense mutations. In all three groups, the C-allele significantly associates with lower p53 expression levels (**Fig. 3D**).

**Figure 3.**
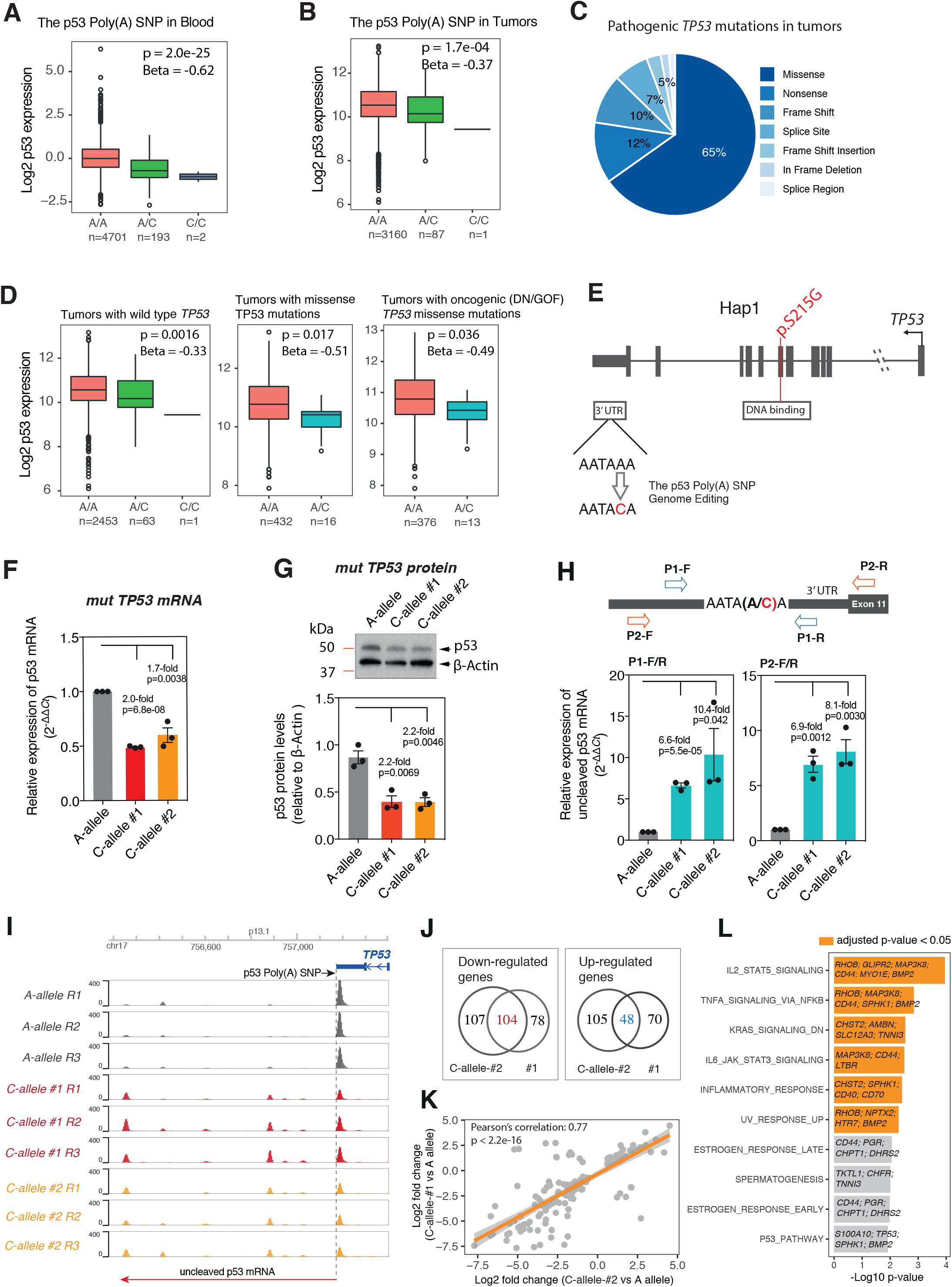
A p53 regulatory cancer risk SNP can affect wild type and mutant p53 in tumors. (A) A box plot of p53 mRNA expression levels on the y-axis (Log2 scale) in blood samples from 4,896 individuals with differing genotypes of the p53 poly(A) SNP (x-axis): 4710 [A/A] homozygotes, 193 [A/C] heterozygotes and 2 [C/C] homozygotes. The central horizontal line indicates the median of each distribution, upper and lower boundaries of the boxes indicate the 3rd and 1st quartiles. The p-value (linear regression) and beta coefficients of the association of the genotype with mRNA levels are depicted. (B) A box plot of p53 mRNA expression levels in 3,248 tumors from individuals with differing genotypes of the p53 poly(A) SNP (x-axis): 3160 [A/A] homozygotes, 87 [A/C] heterozygotes and 1 [C/C] homozygote. The p-values and beta coefficients were determined using a linear model and are displayed above the plots. (C) A pie chart of the percentages of different classes of pathogenic p53 mutations in the 3,985 tumors of the TCGA cohort that had *TP53* sequence information available. (D) Box plots of p53 mRNA expression levels in tumors from individuals with differing genotypes of the p53 poly(A) SNP. The mRNA levels are depicted for individuals with wild type *TP53* (left), missense *TP53* mutations (center) and oncogenic *TP53* missense mutations (right). The p-values and beta coefficients were determined using a linear model. (E) A schematic diagram of the p53 mutational status and CRISPR/cas9-mediated genome editing strategy in Hap1 cells. The somatic p53 mutation in the DNA binding domain, and the poly(A) SNP minor-allele C in 3’-UTR, are highlighted in red. (F) A bar plot of p53 cDNA levels for each genotype in Hap1 cells, measured using qRT-PCR normalized to GAPDH. Error bars represent SEM of 3 independent experiments. p-values are depicted and were calculated using a two-tailed t-test. (G) A bar plot of p53 protein levels for each genotype in Hap1 cells, measured using densitometric analyses of results from Western blot analyses (upper pane) and normalized to β-actin. Error bars represent SEM of 3 independent experiments. p-values were calculated using a two-tailed t-test. (H) A schematic overview of the qRT-PCR strategy to measure the levels of unleaved p53 mRNA in Hap1 cells of differing genotypes (upper). Two bar plots of uncleaved p53 mRNA levels for each genotype in Hap1 cells, measured using qRT-PCR normalized to GAPDH (lower). Two sets of primers (P1-F/R and P2-F/R) were used to amplify the p53 pre-mRNAs. (I) The results of 3’ RNA sequencing of logarithmically growing cells from multiple clones and replicates of cells with the p53 poly(A) SNP C-alleles (red and orange tracks) and with multiple replicates of the A-allele clone (grey bars). The track abundance is plotted for each replicate for the RNAs found at the 3’ end of the *TP53* gene and a diagram of the gene is found above the plots for reference. A vertical dotted line and horizontal arrow indicates the uncleaved p53 RNAs. (J) Venn diagrams of the down-regulated (left) and the up-regulated (right) genes identified in the C-allele-containing clones (#1, right; #2, left) relative to the A-allele-containing clone. (K) A scatterplot showing the Pearson’s correlation between the log2 fold change values of the common significantly differentially expressed genes identified in two edited clones with the C-allele compared to the clone with the A-allele (adjusted p values < 0.05, fold change >1.5). (L) A bar plot of the −log10 p-values for the top 10 enriched pathways amongst the commonly down-regulated 104 genes. The pathways with FDR-adjusted p-values less than 0.05 are indicated in orange. The transcripts in each pathway that were enriched are noted.

To further study the effect of the p53 poly(A) SNP on p53 expression in cancer cells, we developed a primarily isogenic cellular model with the two different alleles in the endogenous p53 locus. Specifically, we utilized Hap1 cells that contain a dominant-negative p53 missense mutation (p.S215G), which results in a mutated DNA-binding domain (34), and which has been found in many cancer types (COSM43951). Using CRISPR/Cas9-mediated genome editing and homologous recombination, we generated clones with either the A-allele or the C-allele (**Fig. 3E** and **Supplementary Fig. 1B**). Consistent with the results found in the TCGA tumors, we found significantly lower p53 mRNA levels in cells with the C-allele relative to the A-allele using qRT-PCR (∼2 fold, p = 6.8e-08 for clone #1 and p = 0.0038 for clone #2; **Fig. 3F**). We also found the C-allele containing cells express less p53 protein: approximately 2-fold (**Fig. 3G**). The impairment of 3’-end processing and subsequent transcription termination by the minor allele of the p53 poly(A) SNP, have been proposed as a mechanism for the genotype-dependent regulatory effects on p53 expression (33). To investigate whether this is also the mechanism by which the C-allele reduces oncogenic mutant p53 levels in cancer cells, we determined the levels of p53 mRNA transcripts not cleaved at the canonical AAUAAA site (uncleaved) relative to the cleaved transcripts using two different approaches. First, using specific probe/primer sets and qRT-PCR, we observed significant 6-10-fold relative enrichments of uncleaved p53 mRNA in cells carrying the C-allele compared to the A-allele (**Fig. 3H**). Next, using data derived from 3’ RNA-sequencing of RNA derived from logarithmically growing cells from multiple clones of each genotype, we also found more uncleaved p53 mRNA in cells carrying the C-allele (red and orange tracks; **Fig. 3I**) relative to A-allele (grey tracks). Together, our data demonstrate that this cancer risk-associated SNP can influence the expression of both wild type and mutant p53 in cancer cells and tumors.

In order to explore whether allelic differences in mutant p53 expression result in allelic-differences in the oncogenic properties of mutant p53 in cancer cells, we next compared the transcriptomes of cells with the different alleles, given the increasing evidence that mutant p53 activities are critical components of oncogenic transcriptional networks (15). We found both C-allele containing clones (less mutant p53) to differentially express a similar number of transcripts relative to the parental cell line (A-allele; more mutant p53; 182 down-regulated and 118 up-regulated genes in clone #1, and 211 down-regulated and 153 up-regulated genes in clone #2; fold change > 1.5, adjusted p value < 0.05; **Fig. 3J**), and the log2 fold change of these differentially expressed genes are highly correlated (Pearson’s r = 0.77; **Fig. 3K**). To examine whether the genotype-dependent transcriptional alterations are associated with changes in mutant p53-associated oncogenic networks, we first excluded the potential clonal effects by selecting genes that are differentially expressed in both C-allele containing clones (104 down-regulated genes and 48 up-regulated genes; **Fig. 3K**; **Supplementary Table S5**). Next, we performed pathway enrichment analyses using the curated Hallmark gene sets (35). We observed the down-regulated genes to be highly enriched in transcripts involved in mutant p53-associated oncogenic networks, such as JAK/STAT, TNF-α/NF-κB and KRAS pathways (**Fig. 3L**; **Supplementary Table S6**). Specifically, these include the *CD44* and *MAP3K8* transcripts, which have been shown to be positively regulated by mutant p53 (36,37). Thus, the p53 poly(A) SNP not only results in allelic-differences in mutant p53 expression, but also in one of its oncogenic properties.

### 4. A p53 regulatory cancer risk SNP associates with patient outcome in a manner that depends on somatic p53 mutational status

*TP53* mutation in tumors has been associated with worse survival or lack of response to therapy in many cancer types (38). Indeed, when we compare patients from the TCGA pan-cancer cohort who have tumors with p53 mutations (2,513), p53 CNV loss (2,655) or both (1,457) to the patients without *TP53* mutation (4,499), loss (4,200) or both (3,168), all three groups displayed shorter progression-free interval (PFI) and worse overall survival (OS) (**Fig. 4A**). To further explore whether p53 regulatory cancer-risk SNPs could have persistent effects on cancer cells and tumors, we next examined whether the p53 poly(A) SNP also associates with allelic differences in clinical outcomes in this pan-cancer cohort. We stratified the cohort into two groups based on p53 somatic alterations and the p53 poly(A) SNP genotypes. We found that in patients with p53 WT tumors, those with the minor C-alleles (less p53 expression; increased cancer risk) have a significantly shorter PFI and worse OS compared to those without the minor alleles (more p53 expression; decreased cancer risk) (**Fig. 4B**, p = 0.0092 for PFI; **Fig. 4C**, p = 0.0059 for OS), but not in patients without stratification (**Supplementary Fig. 1C**). An inverted, but not significant trend, among the patients with somatic *TP53* mutations is noted. The lack of significance is unsurprising do to the low minor allele frequency (**Fig. 4B-C**). Similarly, significant, p53 mutational status-dependent, associations between the p53 poly(A) SNP and PFI can be found when we restrict our analyses to breast cancer patients only (**Fig. 4D**; **Supplementary Fig. 1C**).

**Figure 4.**
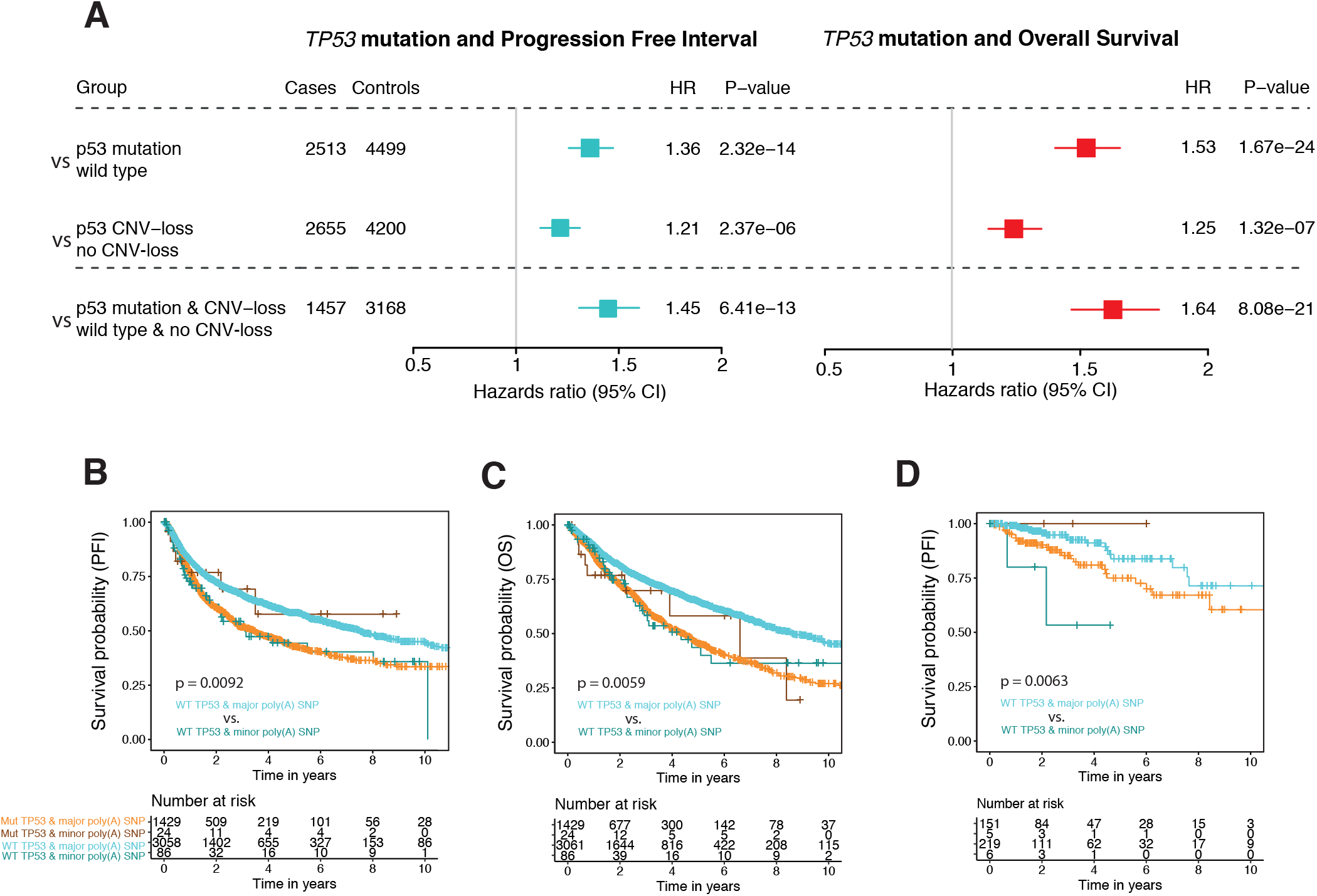
A p53 regulatory cancer risk SNP associates with patient outcome in a manner that depends on somatic p53 mutational status. (A) A forest plot of the progression free intervals (PFI) and overall survival rates (OS) of 7,012 cancer patients (pan-cancer TCGA cohort) stratified by the somatic p53 mutational status: wild type, copy number loss (CNV-loss), and p53 mutation. Number of patients in each group are indicated on the left, and the hazard ratios (HR) comparing PFI and OS in patients with or without mutations are indicated on the right. HR and logrank p values are also displayed and were calculated using Cox proportional hazards model. The error bars represent 95% confidence intervals. (B–C) Kaplan-Meier survival curves for PFI (B) and OS (C) in a total of 4,625 cancer patients carrying either the major or the minor allele of the p53 poly(A) SNP and/or somatic *TP53* mutations. Curves were truncated at 10 years, but the statistical analyses were performed using all of the data (logrank test). Below each plot, the number of patients for each time point, and genotype class, are indicated. (D) Kaplan-Meier survival curves for PFI in a total of 381 breast cancer patients carrying either the major or the minor allele of the p53 poly(A) SNP and/or somatic *TP53* mutations.

### 5. p53 pathway genes with cancer risk SNPs associate with cellular chemosensitivities to p53 activation

Somatic p53 mutation or inhibition is associated with resistance to targeted and DNA damaging chemotherapies and consequently, various therapeutic efforts have been designed around restoring p53 WT activity to improve p53-mediated cell killing (39). The identification of a p53 regulatory cancer risk SNP that affects p53 expression levels, activity, mutational status and tumor progression (as demonstrated for the p53 poly(A) SNP) points to other potential entry points for therapeutically manipulating p53 activities guided by these commonly inherited variants. If true, we reasoned that the p53 pathway genes that harbor cancer risk SNPs could be more likely to associate with differential p53-mediated cancer cell killing relative to other p53 pathway genes. In total, there are 1,133 GWAS implicated cancer-risk SNPs (lead SNPs and proxies) in 41 out of 410 annotated p53 pathway genes (KEGG, BioCarta and PANTHER and/or direct p53 target genes (40)) (**Supplementary Table S7**). The 1,133 SNPs associate with 19 different cancers with an average odds ratio (OR) of 1.17, ranging from 1.03 to 3.07 and with two SNPs associating with significantly larger ORs (**Fig. 5A**): the p53 poly(A) SNP with an odds ratio of up to 2.79 for glioma and SNPs in the p53 target gene KITLG with an odds ratio of up to 3.07 for testicular germ cell tumor risk (TGCT, **Fig. 5A** red dots).

**Figure 5.**
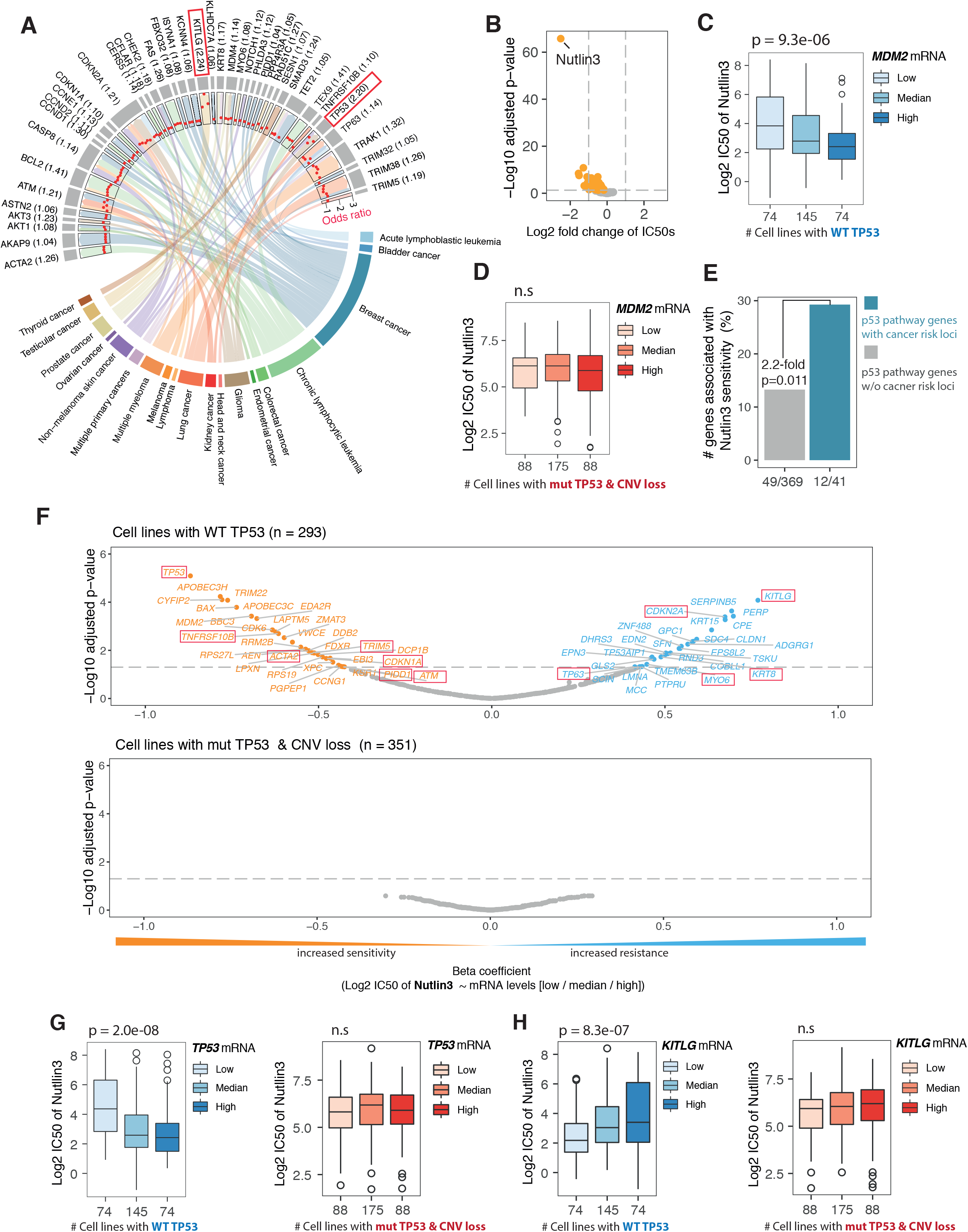
p53 pathway genes with cancer risk SNPs associate with cellular chemosensitivities to p53 activation. (A) A Chord Diagram of 102 cancer GWAS lead SNPs in 41 p53 pathway genes (upper) that associate differential risk to a total of 19 different cancer types (lower). The width of the connecting band indicates the number of lead SNPs for each association. A dot plot of the odds ratios for each association is presented in the inner circle and with red dots. The median odd ratio for each association is presented in parentheses next to the gene name. The two genes, *TP53* and *KITLG* with the highest odds rations are boxed in red. (B) A volcano plot of 304 drugs and their association with differential sensitivity in 311 cancer cell lines carrying WT *TP53* relative to 365 cell lines with *TP53* somatic alterations. −Log10 adjusted p-values (linear regression and FDR-adjusted) are plotted against the Log2 fold change of the average IC50 concentrations (*TP53* WT vs. mutant and CNV loss). The horizontal dashed lines represent the FDR-adjusted p value of 0.05. The 117 drugs significantly associated with differential sensitivity are labeled in orange. (C-D) Box plots of the Log2 average IC50 values of Nutlin3 in cells either with low, medium or high MDM2 mRNA levels and wild type (C) or mutant (D) *TP53*. The number of cell lines analyzed in each group is indicated below the relevant plot. The p-values were determined using a linear model and are displayed above the plots. (E) A bar graph of the percentage of the p53 pathway genes with cancer GWAS loci that also associate with Nutlin3 sensitivity compared with the p53 pathway genes without cancer GWAS risk loci. (F) Two volcano plots of the level of the associations between the transcript levels of the 410 *TP53* pathway genes and Nutlin3 sensitivities in cancer cell lines with either WT (upper) or mutant (lower) *TP53*. The −Log10 adjusted p-value for each association is plotted against the beta coefficient. The horizontal dashed lines represent the FDR-adjusted p value of 0.05. (G-H) Box plots of the Log2 average IC50 values of Nutlin3 in cells either with low, medium or high mRNA (G: *TP53*; H: *KITLG*) levels and wild type *TP53* (left panel in blue) or mutant *TP53* (left panel in red). The p-values were determined using a linear model as described above.

To identify those p53 pathway genes whose expression associates with differential p53-mediated cancer cell killing, we mined a drug sensitivity dataset with both somatic genetic and gene expression data (GDSC, Genomics of Drug Sensitivity in Cancer; 304 drugs across 988 cell lines) (41). Of the 304 drugs analyzed, 127 drugs demonstrated heightened sensitivity in cell lines with WT *TP53* compared to those with *TP53* mutations (adjusted p < 0.05; **Fig. 5B** orange dots; **Supplementary Table S8**). The p53 activator, the direct MDM2 inhibitor (Nutlin3), was the clear outlier, whereby a 5.8-fold greater sensitivity was found in *TP53* wild type cells (adjusted p = 2.6e-67). Moreover, in *TP53* wild type cells, but not *TP53* mutant cells, we found a further increased sensitivity to Nutlin3 in those cells with heightened expression of MDM2 mRNA (**Fig. 5C-D**). In total, 61 of the 410 p53 pathway genes (14.9%) showed similar associations with transcript levels and Nutlin3 sensitivities in *TP53* wild type cells, but not mutant cells (adjusted p < 0.05; **Fig. 5 F**; **Supplementary Table S9**). Interestingly, when we restrict our analysis to those 41 p53 pathway genes with cancer risk SNPs, we note a 2.2 fold enrichment relative to pathway genes without cancer risk SNPs (p=0.011, Fisher’s exact test, **Figure 5E**). Specifically, 12 (29% compared to 13%) showed significant associations between the mRNA expression levels and Nutlin3 sensitivities in TP53 wild type cells, but not mutant cells (adjusted p < 0.05; **Fig. 5 E and 5F** red squares). These observations lend support to the hypothesis that the p53 pathway genes that harbor cancer risk SNPs are more likely to associate with differential p53-mediated cancer cell killing relative to other p53 pathway genes.

For 7 of the cancer risk SNP containing p53 pathway genes, higher expression levels associated with heighted sensitivity towards Nutlin3 (**Fig. 5F** orange dots and red squares), while for 5 of the genes, higher expression levels associated with less sensitivity (**Fig. 5F** blue dots and red squares). For the first group, p53 is clearly the most significant transcript and for the second, KITLG. Specifically, cell lines with wild type *TP53* and more p53 mRNA expression are more sensitive to Nutlin3, as could be expected with a p53 activator (p = 2.0e-08; **Fig. 5G** left panel), while cell lines with wild type *TP53* and more KITLG mRNA expression are less sensitive to Nutlin3 (p = 8.3e-07; **Fig. 5H** left panel). It is important to note that no such association between expression level of these genes and Nutlin3 sensitivities is found in cell lines with mutant p53 (**Fig. 5F** lower panel**; Fig. 5G-H** right panel). *KITLG* (Kit Ligand, also Stem Cell Factor) encodes the ligand for the c-KIT oncogene, which activates a pro-survival signaling cascade that can be inhibited by multiple receptor tyrosine kinase inhibitors (RTKs) used for treatment of multiple cancers (42). In contrast, directly pharmacologically targeting p53 itself has proven challenging during the last three decades.

### 6. The p53-bound cancer risk SNPs in *KITLG* associate with patient outcome

The above-described analysis of p53 pathway genes harboring cancer risk SNPs thus points to KITLG as a promising candidate druggable gene whose heightened expression associates with less p53-mediated cancer cell killing. The identified TGCT risk locus falls within an intron of *KITLG* and contains a polymorphic p53 response element (p53-RE) (43). Somatic amplification of the MDM2 oncogene is inhibitory to p53, leading to pro-survival effects on p53 wild-type cancer cell; *MDM2* is recognized thus to be a targetable entity within the p53 pathway. Hence, we sought to explore whether the p53-dependent up-regulation of *KITLG* expression could lead to similar pro-survival phenotypes in TGCT. To begin to test this, we first had to fine-map the locus for both the association with TGCT risk and p53 occupancy, in order to determine if the greatest association with TCGT risk was indeed found in the genomic region occupied by p53. Using data from the 1,000 Genomes Project as a reference panel, we imputed the genotypes for two independent TGCT GWAS cohorts (44,45). The strongest TGCT GWAS signal lies in intron 1 of *KITLG*, which contains 6 common genetic variants that are in high LD in Europeans (r^2^ >0.95) (red square, **Fig. 6A** and **Supplementary Fig. 2A**), including the 2 lead SNPs (rs3782181 and rs4474514) identified by multiple GWAS studies. Importantly, these clustered SNPs in the *KITLG* gene reside in a region occupied by p53 in 20 of the 30 p53 ChIP-seq datasets analyzed (**Supplementary Table** S10). The cluster spans a region just over 1 kb (1,355 base pairs) (**Fig. 6B**), and contains 4 SNPs (rs7965365, rs3782180, rs4590952, and rs4474514), including the previously identified rs459052, which reside in predicted p53-REs as determined by a position weight matrix (PWM) developed using p53-REs in target genes (43) (**Fig. 6B**, red bars; **Supplementary Table** S11).

**Figure 6.**
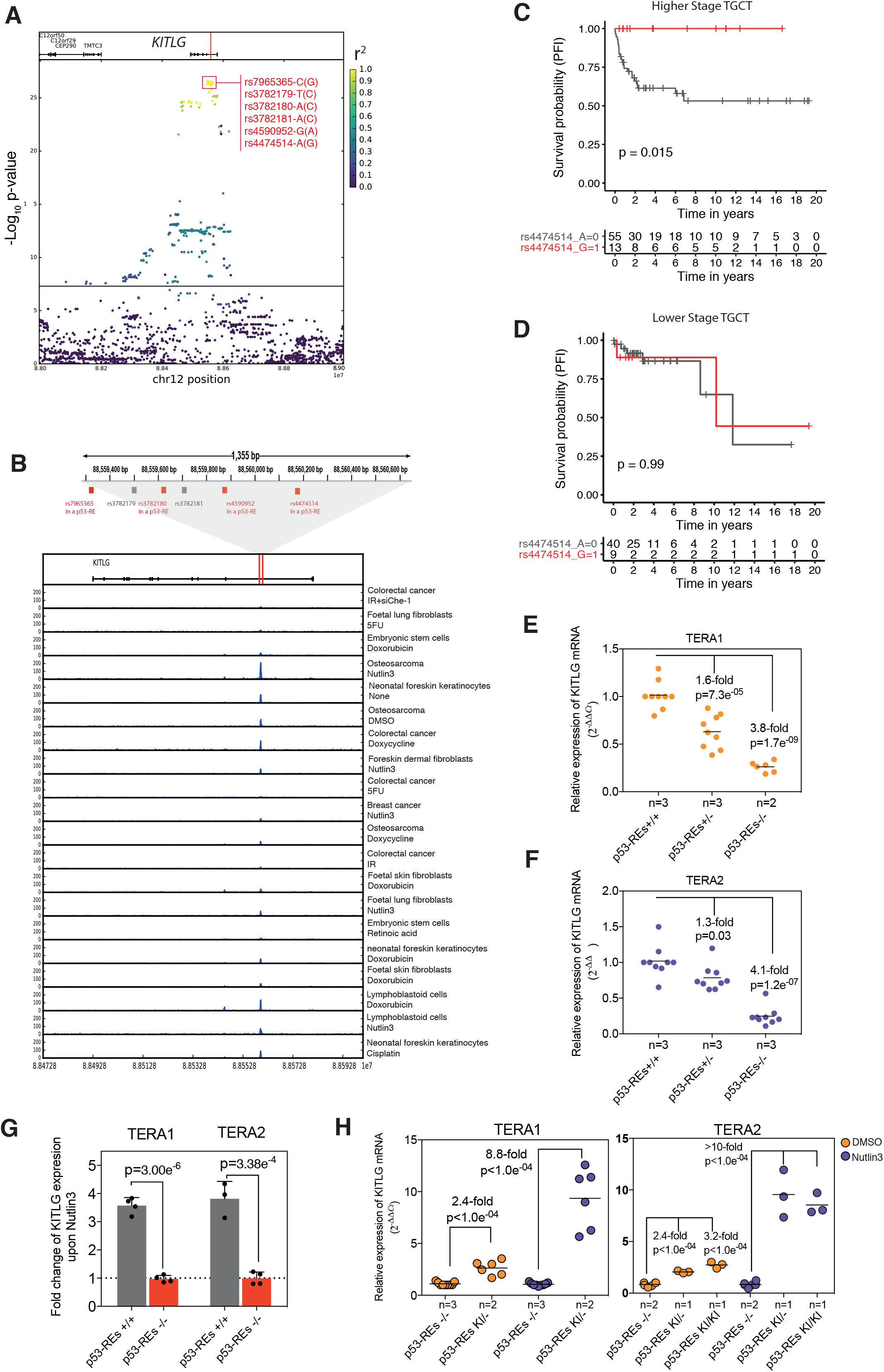
The p53-bound cancer risk SNPs in *KITLG* associate with patient outcome and *KITLG* expression. (A) Genetic fine mapping identified 6 SNPs with the strongest TGCT GWAS signal (high −Log_10_ p-values) and which are in high linkage disequilibrium in Europeans (r^2^ >0.95; red square). The color scale in the right panel indicates the linkage disequilibrium (r^2^) at this locus. (B) A highly p53-occupied risk locus contains four SNPs reside in predicted p53-REs (red boxes). (C-D) Kaplan-Meier survival curves for PFI in high-stage (C: 68 patients) or low-stage TGCT patients (D: 49 patient) carrying either the risk (in grey) or non-risk allele (in red) of the KITLG SNP. p value was calculated using log-rank test. (E-F) *KITLG* gene expression (in TCGT cell lines TERA1 and TERA2, as measured in non-edited clones (p53-REs+/+), heterozygous knock-out clones (p53REs+/−) and homozygous knock-out clones (p53-REs−/−) using qRT-PCR normalized to GAPDH. In total, 2 to 3 clones of each genotype were analyzed in 3 independent biological replicates. Black lines indicate the mean expression; *n*, the number of clones per genotype. p-values were calculated using a one-way ANOVA, followed by Tukey’s multiple comparison test. (G) A bar graph of the fold change in *KITLG* cDNA levels after Nutlin3 treatment, measured using qRT-PCR normalized to GAPDH and a DMSO control. Error bars represent SEM of 2 clones for each genotype and in 2 independent experiments. p-values were calculated using a two-tailed t-test. (H) Dot plots of *KITLG* cDNA levels that were measured using qRT-PCR and normalized to GAPDH. Each dot represents the mean of 3 technical replicates for a given biological replicate.

Next, we explored whether these p53-bound germline TGCT risk-associated SNPs could also have persistent effects on tumors during the course of the disease. To begin to test this, we first evaluated potential associations with disease progression. To do this, we determined the PFI of 118 TGCT patients of European ancestry with p53 WT tumors (TCGA, **Supplementary Table** S12). We grouped the patients with higher stage tumors (IS, II, III) or lower stage (I) TGCT (**Supplementary Fig.** 2B) and found that the cancer risk SNP(s) associated significantly with PFI in patients with higher stage tumors, whereby the alleles associated with greater TCGT risk (better predicted p53 binding) associated with shorter PFI (p = 0.015; **Fig. 6C-D**).

### 7. The p53-bound cancer risk region is a p53-regulated KITLG enhancer in cancer cells

We next tested whether this TGCT risk locus remained a p53-regulated enhancer in cancer cells. To do this, we deleted the 1-kb region from two testicular germ cell tumor-derived cell lines (TERA1 and TERA2) with WT p53 and homozygous for the p53-bound TGCT risk alleles (p53-REs+/+) (**Supplementary Fig. S3A-C**). In all clones tested (at least 2 clones for both the non-edited, the heterozygous KO and the homozygous KO cells), we found significantly higher *KITLG* RNA levels in non-edited p53-REs+/+ clones, compared to either the heterozygous KOs p53-REs+/− clones (an average of 1.6 fold for TERA1, p = 7.3e-05; 1.3 fold for TERA2 cells, p = 0.03) or the homozygous KOs REs−/− clones (an average of 3.8 fold for TERA1, p = 1.7e-09; 4.1 fold for TERA2 cells, p = 1.2e-07; **Fig. 6E-F**). We then treated TERA1 and TERA2 p53-REs+/+ cells with the p53-activating agent Nutlin3 (an MDM2 inhibitor) and observed ∼4-fold induction of *KITLG* over DMSO treated cells in both cell lines (**Fig. 6G**, grey bars). Treatment of the p53-REs−/− clones with Nutlin3 showed no measurable induction of *KITLG* (**Fig. 6G**, red bars versus grey bars). Moreover, the transcripts from genes that lie approximately 2 Mbp on either side of *KITLG* were measured in the clones of both genotypes, but no significant differences were found between the p53-REs−/− and p53-REs+/+ clones (**Supplementary Fig. 4A**). We also tested the dependency of the p53-bound enhancer on *KITLG* expression and/or induction by reinserting it into the p53-REs−/− clones (**Supplementary Fig.** S3G-H). Re-integration rescued basal expression, resulting in significantly higher *KITLG* RNA levels in the knock-in (KI) clones of both cell lines relative to the p53-REs−/− (**Fig. 6H**). The KI clones also rescued the p53-dependent induction of *KITLG* expression relative to the p53-REs−/− (**Fig. 6H**).

To evaluate whether the TGCT risk haplotype in *KITLG* affects this enhancer activity in TGCT, we compared the endogenous enhancer activities of the risk haplotype and non-risk haplotype in two other TGCT cell lines (Susa-CR and GH) that we engineered to be heterozygous for this locus (**Supplementary Fig. S3D-F**). We assessed *KITLG* levels in the non-risk haplotype (p53-REs-/non-risk) and the risk haplotype (p53-REs-/risk). At basal levels, we found significantly higher *KITLG* expression in non-edited p53-REs+/+ clones compared to the p53-REs-/non-risk clones (**Supplementary Fig. 4B-C**). When we treated these multiple clones with Nutlin3 to activate p53, we observed significant higher *KITLG* expression in-/risk relative to-/non-risk clones (**Supplementary Fig. 4B-C**), indicating a gain of p53-mediated enhancer activity in association with the risk haplotype. Together, these data demonstrate that the p53-bound region associated with TGCT risk and progression is a p53-regulated enhancer for *KITLG* expression in TGCT cancer cells.

### 8. p53/*KITLG* pro-survival signaling can attenuate responses to p53-activating agents

As mentioned above, somatic amplification of the MDM2 oncogene results in pro-survival phenotypes in p53 wild type cancer cells, thus making it an attractive drug target to increase p53-mediated cancer cell killing. Thus, we next explored whether the p53-dependent up-regulation of *KITLG* expression results in similar pro-survival phenotypes in TGCT cells. *KITLG* acts through the c-KIT receptor tyrosine kinase to promote cell survival (42), so first we knocked down c-KIT expression in TGCT cells and measured cell proliferation and migration rates. Reduced c-KIT expression in TERA1 and TERA2 cells substantially attenuated proliferation and migration, supporting c-KIT-dependent pro-survival activity in TCGT (**Supplementary Fig. S5A-B**). Next, to explore if the p53-mediated up-regulation of *KITLG* has a similar pro-survival effect on TGCT cells, we compared proliferation and migration rates of the p53-REs+/+ clones (more *KITLG*) relative to the p53-REs−/− clones (less *KITLG*). Consistent with the relative reduction in *KITLG* expression levels and the effects of c-KIT knock-down on TGCT proliferation and migration, p53REs−/− clones grew and migrated significantly more slowly than p53-REs+/+ clones (**Supplementary Fig. S5C-D**).

These results link p53 driven *KITLG*/c-KIT signaling with oncogenic pro-survival phenotypes in TGCT, such as heightened proliferation and migration. To determine the impact *KITLG*/c-KIT has on cellular sensitivities to p53-activating therapies we used cells with reduced c-KIT expression and treated then with Nutlin3. c-KIT knock-down resulted in a 2-fold increased sensitivity to Nultlin3, and increased levels of cleaved caspase3, relative to control cells (**Supplementary Fig. S6A-B**). These data suggest that c-KIT signaling can attenuate cellular chemosensitivities to p53-activating therapies. To explore if p53-mediated up-regulation of *KITLG* has a similar effect we measured IC50 values for Nutlin3 in p53-REs +/+, p53-REs −/− and p53RE KI clones for both TERA1 and TERA2 cells, and observed a significant reduction in IC50 values in the p53-REs−/− cells relative to p53-REs+/+ cells upon Nutlin3 treatment (TERA1: 3.0-fold, p = 0.021; TERA2: 1.8-fold, p = 7.1e-04; **Fig. 7A**). We were able to rescue the increased Nutlin3 sensitivity of p53RE−/− clones in KI cells (TERA1: 2.2-fold, p = 0.035; TERA2: 1.5-fold, p = 0.033; **Fig. 7A**). Consistent with these observations, we saw increases in cleaved Caspase3 and cleaved PARP1 levels in p53-REs−/− cells relative to p53-REs+/+ cells (**Supplementary Fig. S6C**), but not in the KI cells (**Supplementary Fig. S6D**). To further test the p53-dependence of these effects, we reduced p53 expression levels and observed reduced expression of cleaved caspase3 after Nutlin3 treatment (**Supplementary Fig. S6E**), and an overall insensitivity towards Nutlin3 in both p53-REs+/+ and p53-REs−/− cells (**Supplementary Fig. S6F**). Thus, *KITLG*/c-KIT signaling promotes cell survival and attenuates cellular chemosensitivities towards a p53-activating agent, and these regulations involve the risk locus in *KITLG*.

**Figure 7.**
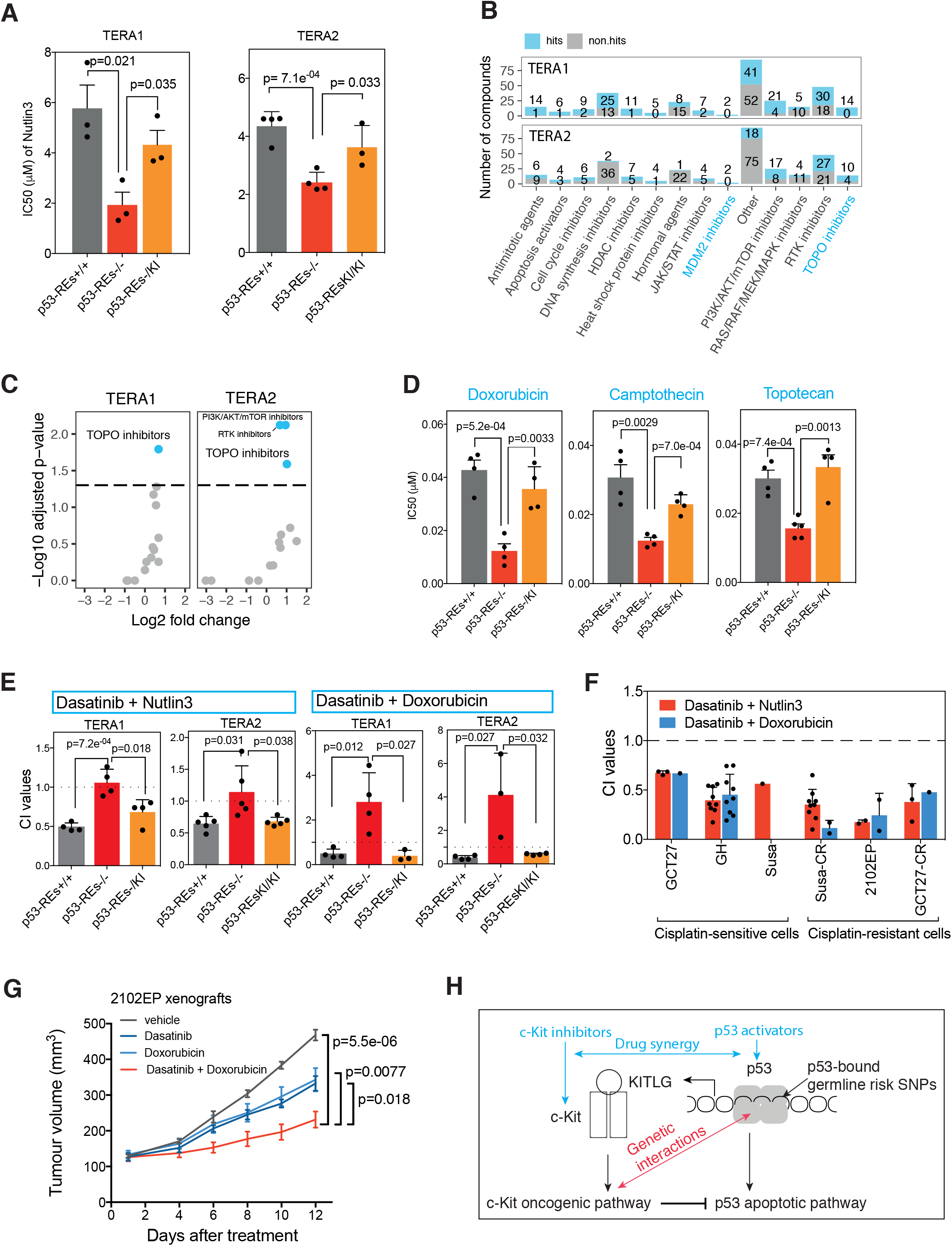
p53/*KITLG* pro-survival signaling can attenuate responses to p53-activating agents. (A) Bar blots of the IC50 values for Nutlin3 of TERA1 and TERA 2 cells. p-values were calculated using a two-tailed t-test and error bars represent SEM in at least 3 independent biological replicates. (B) Bar plots depicting the number of hits and “non-hits” for each of the 14 drug classes examined. (C) Scatter plots of the fold enrichment of hits on the x-axis (log_2_ scale), and the adjusted p-value on the y-axis (−log_10_ scale), amongst each drug class relative to the total compounds in 14 drug classes. The horizontal dashed lines represent the FDR-adjusted p value of 0.05. (D) Bar plots of average IC50 values for 3 TOPO inhibitors in p53-REs+/+ (grey bars, two clones), p53-REs−/− (red bars, two clones) p53-REs-/KI (orange bars, one clone) of TERA1 cells. Error bars represent SEM of at least two independent biological replicates. (E) Bar plots of combination indexes of Dasatinib with Nutllin3 or Doxorubincin in p53-REs+/+ (grey bars, two clones), p53-REs−/− (red bars, two clones) and knock-in clones (orange bars, one clone) of TERA1 and TERA2 cells. (F) Bar plots of combination indexes of Dasatinib with Nutllin3 or Doxorubincin in panel of TGCT cell lines. (G) Growth curves of 2102EP xenograft tumors treated with vehicle, Doxorubicin, Dasatinib or the combination of Doxorubicin and Dasatinib. Error bars represent means ± SEM (n=6). (H) A diagram depicting the development of more effective therapy combinations by modulating both the cell death and survival functions of p53 based on both the inherited and somatic genetics of the patient.

The synthetic viable interaction between *KITLG* and p53 activation by Nutlin3 in TGCT cancer cells suggests *KITLG* should show similar synthetically viable interactions with chemotherapeutic agents which lead to DNA damage, given the role of p53 in responses to DNA damage (9,46). To test this idea, we utilized one p53-REs+/+ and one p53-REs−/− clone of both TERA1 and TERA2, and screened 317 anti-cancer compounds to identify agents that, like Nutlin3, kill significantly more cells at lower concentrations in p53-RE−/− clones than in p53+/+ clones (**Supplementary Fig. S7A**). The screen was performed in duplicate. The Pearson Correlation Coefficient, a measurement for inter-assay variability, averaged 0.98 and an average Z-factor, a measure employed in high throughput screens to measure effect size, of 0.69 for all plates was recorded, leading to high confidence in the primary screen positive hits (**Supplementary Table** S13). We identified 198 compounds in the TERA1 screen and 112 compounds in the TERA2 screen that showed heightened sensitivity in p53-RE−/− cells in at least one of the 4 different concentrations tested (≥1.5 fold in both replicates; **Supplementary Fig. S7B**, blue dots). One hundred of these agents overlapped between TERA1 and TERA2 (1.7-fold, p = 1.1e-21; **Supplementary Fig. S7B**, Venn diagram), suggesting a potential shared mechanism underling the differential sensitivities. These 100 agents can be classified into 14 different compound classes (**Fig. 7B**; **Supplementary Table** S14). Consistent with our previous results, two *MDM2* inhibitors in the panel of compounds, Nutlin3 and Serdemetan, were among the 100 overlapping agents (**Fig. 7B**).

In TERA1, the 198 compounds were significantly enriched in topoisomerase inhibitors after correction for multiple hypothesis testing (**Fig. 7C**, left panel). In TERA2, the 112 compounds were also significantly enriched in topoisomerase inhibitors, but also in PI3K/AKT/mTOR inhibitors and receptor tyrosine kinase (RTK) inhibitors (**Fig. 7C**, right panel). We found a significant and consistent enrichment of topoisomerase inhibitors in both cell lines (14 compounds in TERA1 [100%] and 10 compounds in TERA2 [71%] of 14 Topo inhibitors screened; **Fig. 7B-C**). Topoisomerase inhibitors induce DNA damage and p53 activation (46,47). To validate the genotype-specific effects of the topoisomerase inhibitors, we determined the IC50 values of three of them, Doxorubicin, Camptothecin. and Topotecan, using MTT measurements in multiple clones of TERA1 cells with differing genotypes. All three agents showed a significant reduction of IC50 values in the p53-REs−/− clones relative to the p53-REs+/+ clones (**Fig. 7D**, grey bars versus red bars). We were able to rescue this increased sensitivity to topoisomerase inhibitors in the p53RE−/− clones in KI cells (**Fig. 7D**, orange versus red bars). Together, these results demonstrate a synthetically viable interaction between the germline risk locus and multiple p53-activating agents that lead to DNA damage.

### 9. Inhibition of c-KIT signaling and p53 activation interact to kill treatment resistant cancer cells

There are many RTK inhibitors that are current therapeutic agents which inhibit c-KIT activity (48). If p53-mediated *KITLG*-dependent pro-survival signaling can attenuate chemosensitivity to p53-activating agents, RTK inhibitors should be able to interact synergistically with p53-activating agents to kill TGCT cells. We therefore tested which RTK inhibitor (known to inhibit c-KIT) kills TCGT cells most efficiently. Of the five FDA-approved RTKs analyzed, Pazopanib, Imatinib, Nilotinib, Suntinib and Dasatinib, the most potent was Dasatinib (**Supplementary Fig. S7C**). To determine potential synergy of RTKs with Nutlin3 in TGCT, we treated TERA1 and TERA2 cells with Dasatinib, and quantitated potential drug-drug interactions by calculating Combination Indices (CI). We observed clear synergistic interactions (CI <1) between Nutlin3 and Dasatinib in both TERA1 and TERA2 p53-REs+/+ cells (**Fig. 7E**, grey bars). These results further support an inhibitory role for p53/*KITLG* pro-survival signaling in cellular responses to p53-activating agents.

To more directly test whether or not the synergistic interaction between Dasatanib and Nutlin3 is mediated by the p53-dependent up-regulation of *KITLG*, we determined the CI values in TERA1 and TERA2 p53-REs−/− cells, wherein p53 cannot induce *KITLG* expression after p53 activation upon Nutlin3 treatment as shown in **Fig. 6G**. Consistent with the requirement of the p53-dependent activation of *KITLG*, no synergy between Dasatanib and Nutlin3 was detected in p53-REs−/− cells (**Fig. 7E**, red bars).

To further investigate if c-KIT inhibition can interact synergistically with p53-activating agents to kill TGCT cells, we explored the interaction between Dasatinib and multiple DNA-damaging chemotherapeutics known to activate p53. We focused on the 3 topoisomerase inhibitors (Doxorubicin, Camptothecin and Topotecan), as well as Cisplatin, a chemotherapeutic agent used to treat TGCT, and which induces DNA damage and p53. In both TERA1 and TERA2, Dasatinib demonstrated significant levels of synergy with each of the DNA-damaging agents tested in p53-REs+/+ cells (**Supplementary Fig. S7D-E**). Similar to Nutlin3, no synergy was detected in p53-REs−/− cells of either cell lines for any combination of agents (**Supplementary Fig. S7D-E**). Furthermore, the synergistic interaction between Dasatinib and the p53-activating agents Nutlin3 and Doxorubin could be rescued by knocking in the p53-bound region in *KITLG* (**Fig. 7E**, orange bars).

As our results thus far were limited to TERA1 and TERA2 cells, we explored potential interactions in four additional TGCT cell lines with wild-type p53 and at least one copy of the haplotype containing the *KITLG* risk allele SNPs; GH (risk/non-risk), Susa (risk/non-risk), 2102EP (risk/risk) and GCT27 (risk/risk). Consistent with the observations in TERA1 and TERA2 cells, Dasatinib synergistically interacted with Nutlin3 across all the cell lines (**Fig. 7F**, red bars) and also with Doxorubicin (**Fig. 7F**, blue bars). Together, these data indicate that *KITLG*/c-KIT pro-survival signaling can attenuate chemosensitivity to p53-activating agents in TGCT and that this attenuation is dependent on the p53-regulated *KITLG* enhancer lying within the germline TGCT-risk locus.

Thus, a more effective therapeutic strategy for TGCT patients could be to modulate both the cell death and cell survival functions of p53, through co-inhibition of p53/*KITLG*-mediated pro-survival signaling together with the co-activation of p53-mediated anti-survival signaling. Such a therapeutic combination could provide an alternative for patients with treatment-resistant disease (49). To investigate this idea, we explored synergistic interactions between c-KIT inhibitor Dasatinib and p53 activators in cisplatin-resistant clones of GCT27 (GCT27-CR) and Susa (Susa-CR) (50), as well as in the intrinsically cisplatin-resistant TGCT cell line 2102EP (51). Similar to the observations in the cisplatin-sensitive TGCT cell lines, Dasatinib and Doxorubicin interacted synergistically to kill all three cisplatin-resistant clones and cell lines (**Fig. 7F**). To determine if the combination treatment could show a greater efficacy in treating tumors, we generated a subcutaneous xenograft model using the 2102EP cell line. Doxorubicin and Dasatinib were given either alone or in combination. Consistent with the observations made in cell culture, treatment of mice engrafted with 2102EP cells revealed stronger anti-tumoral effects with the Dasatinib/Doxorubicin pair relative to single drug treatments (p = 0.0077 versus the Dasatinib group, and p = 0.018 versus the Doxorubicin group; **Fig. 7G**). This dosing regimen was well tolerated with no body weight loss in mice (**Supplementary Fig. S7F**).

## Discussion

Cancer therapies targeting somatic mutations are associated with variable responses, eventual high failure rates and the development of drug resistance. Somatic genetic heterogeneity among tumors is a major factor contributing to differences in disease progression and therapeutic response (1). In this study, we demonstrate that germline cancer-risk SNPs could influence cancer progression and potentially provide information guiding precision medicine therapy decisions. Our approach focused on cancer-risk SNPs in the p53 signaling pathway and provided evidence that they can have persistent effects on tumors in regards to p53 mutational status, gene expression, cellular signaling, progression and chemo-sensitivity. First, we demonstrated that cancer risk SNPs in the p53 pathway genes can influence whether or not a tumor contains a somatically mutated *TP53* locus (**Fig. 1**–**2**). We demonstrated that the cancer risk SNP, the p53 poly(A) SNP rs78378222 affects the expression of both wild-type and mutant p53 in tumors and interacts with p53 somatic mutational status to modify both cancer susceptibility and progression (**Fig. 1**–**4**). We went on to demonstrate that p53 pathway genes that harbour cancer risk SNPs, as a whole, are more likely to associate with differential p53-mediated cancer cell killing relative to other p53 pathway genes. More specifically for *KITLG,* we demonstrated that the risk alleles of the TCGT-associated SNPs result in the p53-dependent increased expression of the pro-survival target gene and can lead to an attenuation of p53-mediated responses to genotoxic therapies, as well as faster progression (**Fig. 5**–**7**). Finally, we determined that, when the pro-survival signal is inhibited, there is more effective p53-mediated cancer cell killing (**Fig. 7**). Our observations illustrate how cancer susceptibility loci can interact with cancer driver genes to influence cancer cell behaviors, cancer progression, identify novel drug-drug interactions and direct molecularly-informed on-targeted combinatorial therapies.

The p53 stress response pathway inhibits cell survival, mediating both tumor suppression and cellular responses to many cancer therapeutics (52). p53 also targets pro-survival genes. Activation of these genes in tumors retaining wild-type p53 provide a survival advantage (53). For example, the p53 target gene, *TIGAR*, which protects cells from DNA damage-induced reactive oxygen species (ROS) and apoptosis, promotes tumorigenesis in a mouse model of intestinal adenoma. We provide human genetic evidence that also supports a tumor-promoting role of p53 pro-survival activities and, in the case of the TGCT risk locus, points to the development of more effective therapy combinations through the inhibition of these pro-survival activities in tumors that retain p53 activity. Less than 1% of TGCTs from the TCGA cohort have a mutated p53 gene. Although TGCTs are one of the most curable solid tumors, men diagnosed with metastatic TGCT develop platinum resistant disease and die at an average age of 32 years (49). There have been few new treatments developed in the last two decades, and current therapeutic approaches can, importantly in context of a cancer of young men, result in significant survivorship issues, including sustained morbidities and delayed major sequelae (49,54). There is a need for more effective treatments with fewer side effects, to improve the survival and quality of life of these patients. Our observations suggest the TGCT *KITLG* risk allele in the polymorphic p53 enhancer leads to increased p53-dependent activation of the pro-survival target gene, *KITLG*, which increases TGCT survival rather than senescence/apoptosis in the presence of active p53 (**Fig. 7**). We demonstrate that co-inhibition of c-KIT and p53 activation interact synergistically to kill platinum-resistant TGCTs with a drug combination (Dasatinib and Doxorubicin) that had limited toxicity in a Phase II clinical trial (55) (**Fig. 7**), suggesting that an effective therapeutic strategy for treatment-resistant TGCTs could be to modulate both the cell-death and cell-survival functions of WT p53 cancers.

Heritable genetic variants can influence the evolution of cancer genomes in patients (3,4), potentially through altered tissue mutation rates, heightened global genome instability (56), or heightened specific mutational processes, for example via inherited variants in pathways such as BRAC1/2, MMR, and the APOBEC3 gene cluster (57). Understanding in BRCA1/2 mutation carriers of the interactions between the inherited variants and somatic genomes of the cancer has already led to better, more personalized treatment options for BRCA1/2 mutation carriers with PARP inhibitors. Here we provide evidence that this could also be extended to the more frequently inherited cancer risk variants identified in GWAS. We demonstrated that cancer-risk p53 pathway SNPs and p53 mutational status can interact to affect tumors in a way that offers potential therapeutic insights. *MDM2* amplification and p53 mutation show a mutual exclusivity in somatic cancer genomes of soft tissue sarcomas, osteosarcomas and glioblastoma, which may extend to other cancer types (58), suggesting that the amplification and over-expression of this p53 inhibitor reduces the necessity of cancers to mutate p53. Support of this hypothesis comes from a study where p53 was preferentially mutated in murine B-cell lymphomas that had been engineered to express lower *MDM2* levels (59). We show that the up-regulation of a pro-survival p53 target gene associates with increased risk for TGCT that rarely mutates p53, which supports the idea that inherited genetic variants could also reduce the necessity of cancers to mutate p53 by increasing the pro-survival/pro-tumor activities of wild-type p53. This hypothesis points to the development of more effective therapy combinations in tumors that retain p53 activity through the inhibition of pro-survival activities, as our work on the *KITLG* locus in TGCT suggests.

Unlike other tumor suppressors, complete loss of p53 activity is not a requirement for cancer initiation. Reduction of p53 activity below a critical threshold is apparently necessary and sufficient for cancer development (60). Another attribute of p53 cancer genetics is the abundance of missense driver mutations relative to simple deletions. These missense mutations may benefit cancers not simply through loss of p53 function, but also through dominant-negative and gain-of-function activities (61), which may include inhibition of p53 expression, or its ability to heterodimerize with wild-type p53, thereby affecting DNA binding and transcriptional regulation. Described gain-of-function activities often include novel interactions with transcription factors and chromatin-bound protein complexes (8). In mice, knock-in p53 gain-of-function mutants displayed a more diverse set of, and more highly metastatic tumors than p53 knock-out mutants (13,14). Many of the factors that regulate wild-type p53 tumor suppression can also regulate mutant p53, including its pro-cancer activities. For example, wild-type p53 mice that express lower levels of *MDM2* show increased p53 levels, a better p53 stress response, and greater tumor suppression, resulting in later and reduced tumor onset in many tissue types. Mutant p53 levels are also increased in these murine models, but cancers are found to arise earlier and harbor gain-of-function metastatic phenotypes (20).

Our SNP associations with inverted cancer risk and somatic p53 mutational status in humans reveal a similar scenario. Specifically, we demonstrated that the C-allele of the p53 poly(A) SNP which can lead to decreased WT and mutant p53 levels in tumors (**Fig. 3**), associates with an increased risk of wild-type p53 cancers, but decreased risk of sub-types with primarily mutant p53 (**Fig. 2**). For example, women with the minor allele associated with an increased risk for the more p53 wild-type breast and ovarian subtypes and a decreased risk for the more mutant subtypes. Together, these observations support a role for germline p53 pathway SNPs not only modulating risk of disease and tumor biology in p53 WT cancers but also in p53 mutant cancers, wherein alleles that increase mutant p53 levels would also increase its pro-cancer activities.

## Methods

### Analysis of oncogenic *TP53* missense mutations in breast and ovarian cancers

We curated *TP53* pathogenic missense mutations by integrating up-to-date functional evidence from both literature and databases. Specifically, we combined the 2 lists of *TP53* driver mutations in human tumors (62,63) to obtain a list of 323 *TP53* driver mutations. To determine which of these 323 *TP53* diver mutations are oncogenic (either dominant negative or gain of function), we relied on two sources of annotations:188 missense mutations were curated to be oncogenic in IARC *TP53* Database (release 18) (64); 1101 missense mutations were ascertained by human cancer cell-based saturation mutagenesis screen (65) (filter criteria: A549_p53WT_Nutlin-3_Z-score > 1 and A549_p53NULL_Nutlin-3_Z-score > 1 and A549_p53NULL_Etoposide_Z-score < −1). In total, we were able to find 218 out of 323 *TP53* pathogenic mutations are oncogenic (**Supplementary Table** S16).

2,262 *TP53* mutations in 2,201 unique breast cancer samples (from 12 studies; exclude 737 duplicate mutations in samples sequenced by multiple studies) and 492 *TP53* mutations in 471 unique ovarian cancer samples (from 3 studies; exclude 477 duplicate mutations in samples sequenced by multiple studies) were downloaded from cBioPortal on 2018-09-14 (http://www.cbioportal.org). All *TP53* missense mutations were extracted and matched with the curated lists of pathogenic and oncogenic *TP53* missense mutations as described above. Then cancers with pathogenic missense mutations and oncogenic missense mutations were counted. Specifically, 1113 out of 2262 (49.2%) *TP53* mutations in breast cancer are pathogenic missense mutations, of which 1012 (90.9%) are oncogenic. Similarly, 260 out of 492 (52.8%) *TP53* mutations in ovarian cancer are pathogenic missense mutations, of which 228 (87.7%) are oncogenic.

### Analysis for subtype heterogeneity SNPs with Breast and Ovarian cancer association studies

Summary statistics of GWASs for breast cancer susceptibility were downloaded on 2018-03-12 (http://bcac.ccge.medschl.cam.ac.uk/bcacdata/oncoarray/gwas-icogs-and-oncoarray-summary-results/), which included summary statistics from case-control association analyses for ER-positive breast cancer cases (ER+BC) and ER-negative breast cancer cases (ER−BC) compared against disease-free controls. Summary statistics of GWASs for ovarian cancer susceptibility were downloaded on 2018-04-16 (https://www.ebi.ac.uk/gwas/downloads/summary-statistics), which included summary statistics for SNP association with low grade serous ovarian cancer (LGSOC), and with high grade serous ovarian cancer (HGSOC). Estimates of effect sizes [log(OR)s] for subtype-specific case-control studies and their corresponding standard errors were utilized for meta-and heterogeneity-analyses using METAL (2011-03-25 release) (66), under an inverse variance fixed-effect model. Cochran’s Q statistic was calculated to test for heterogeneity and the I^2^ statistic to quantify the proportion of the total variation that was caused by heterogeneity.

### Assigning p53 mutational status to TCGA tumour samples and the association testing

The p53 gene mutation profiles in TCGA primary tumors were downloaded from the TCGA data portal (https://gdc-portal.nci.nih.gov/). These p53 mutation calls (1,245 unique mutations in 3,956 tumors) were classified into pathogenic (1,097 unique mutations in 3,895 tumors), benign (143 unique mutations in 148 tumors), or unclear (5 unique mutations in 5 tumors) based on curated datasets (63,64). The p53 pathogenic missense mutations were further annotated as loss of function, or oncogenic (either dominant negative or gain of function) as described above. Tumors without p53 mutations were assigned as p53 WT; Tumors with at least one pathogenic p53 mutations were assigned as p53 mutant; Tumors with only benign and/or unclear p53 mutations were assigned as p53 benign/unclear; Tumors with only pathogenic p53 missense mutations were assigned as p53 missense mutant; Tumors with only oncogenic p53 missense mutations were assigned as p53 oncogenic missense mutant. The copy number profiles of *TP53* in TCGA primary tumors were retrieved from the Broad GDAC Firehose (https://gdac.broadinstitute.org/) through the fbget tool (v0.1.11 released Oct 31 2017). The association testing was performed using a two-sided Fisher exact test with PLINK (67).

### Cancer GWAS SNPs

The GWAS catalog was downloaded on 2018-02-28 (https://www.ebi.ac.uk/gwas/). We selected the GWAS significant lead SNPs (p-value <5e-08) in Europeans, and retrieved the associated proxy SNPs using the 1000 Genomes phase 3 data through the web server: rAggr (http://raggr.usc.edu). In brief, we selected the GWAS lead SNPs that were identified in European ancestry cohorts, and only defined proxies that met the following criteria: Population: EUR; Min MAF: ≥ 0.01; R2 range: ≥ 0.8; Max distance: 500KB; Max # Mendel error: 1; HWE p-value: 1e-6; Min genotype %: 95. All proxies were mapped to the Ensembl Release 91 (dbSNP build 150) to retrieve the hg38 genomic coordinates using R package biomart. In total, we retrieved a total of 283,240 GWAS SNPs. Next, we isolated the 28,592 cancer GWAS SNPs, including 1,225 lead SNPs and 27,367 proxies, by mapping the GWAS SNPs to 106 unique cancer traits that are distributed into 27 distinct cancer types.

### Pathway enrichment analysis

The pathway gene sets of KEGG and Hallmark were downloaded from the Molecular Signatures Database (http://software.broadinstitute.org/gsea/msigdb/index.jsp). The known p53 direct target genes were downloaded from (40). cis-eQTL datasets were obtained form GTEX (https://gtexportal.org/home/datasets; V7 and qval ≤ 0.05), NESDA/NTR (https://eqtl.onderzoek.io/index.php?page=download) and PancanQTL (http://bioinfo.life.hust.edu.cn/PancanQTL/download).

The hypergeometric distribution enrichment analysis was performed as described in (6). Significance was determined using PHYPER function as implemented in R and multiple hypotheses testing by Benjamini-Hochberg correction.

### RNA-seq analysis

3’ RNA-seq library was prepared using a standardised protocol followed by sequencing using a HiSeq4000 platform (Illumina) at the Oxford Genomics Centre (Wellcome Trust Centre for Human Genetics, Oxford, UK). Sequencing reads were mapped to hg19 using the HISAT2 alignment algorithm (version 2.1.0). The aligned Binary-sequence Alignment Format (BAM) files were used to determine the transcript counts through featureCounts (version 1.6.2). For differential expression analysis, the raw read counts were used as input into the R package DESeq2 (version 1.24.0) for analysis.

### eQTL analysis in normal tissue and TCGA tumors

Data for the eQTL analysis of rs78378222 in normal human tissue are from two studies: the Netherlands Study of Depression and Anxiety (NESDA) and the Netherlands Twin Register (NTR) that consisted of 4,896 blood samples with European ancestry (22). Data for the eQTL analysis of rs78378222 in human tumors were obtained from TCGA (68). The p53 gene expression profiles in TCGA primary tumors were retrieved from the Broad GDAC Firehose (https://gdac.broadinstitute.org/) through the fbget tool (v0.1.11 released Oct 31 2017). eQTL effects were determined with a linear model approach with p53 mRNA expression level as dependent variable and SNP genotype values as independent variable.

### Genotype imputation and population stratification

Genotype data was obtained and filtered as described in (3). Briefly, we obtained genotype calls from the Birdsuite-processed (69) Affymetrix 6.0 SNP arrays for matched normal samples from the TCGA data portal (https://gdc-portal.nci.nih.gov/), set low confidence SNP calls to missing, filtered individuals and SNPs with < 95% call rate and SNPs with MAF < 1% and imputed untyped genotypes using the secure Michigan Imputation Server (70). We used a PCA analysis over genotypes to remove samples that did not cluster tightly with Europeans from the HapMap III reference population.

### TCGA survival analysis

TCGA clinical data was downloaded from recently updated Pan-Cancer Clinical Data Resource (TCGA-CDR) (71). Overall survival (OS) and progression-free interval (PFI), the two most accurate clinical outcomes using the current TCGA data, were added to primary tumors. Of the 7,021 TCGA patients that are clustered tightly with Europeans, OS and PFI data was available for 6,979 and 6,977 patients, respectively. A Cox proportional hazards regression model was used to calculate the hazard ratio, the 95% confidence interval and p values for two-group comparisons. The log-rank test was used to compare the difference of Kaplan-Meier survival curves.

### GDSC drug sensitivity analysis

*TP53* mutation, copy number, mRNA expression data, and drug IC50 values for the cancer cell lines were downloaded from Genomics of Drug Sensitivity in Cancer (GDSC; release-8.1). Specifically, a list of the mutated genes “mutations_20191101.csv”, the processed CNV data “cnv_gistic_20191101.csv” and RNAseq gene expression data “rnaseq_read_count_20191101.csv” were downloaded from https://cellmodelpassports.sanger.ac.uk/downloads. The drug response data (GDSC1_fitted_dose_response_15Oct19) was downloaded from https://www.cancerrxgene.org/downloads/bulk_download.

Cell lines without p53 mutations were assigned as p53 WT; Cell lines with *TP53* somatic mutations and copy-number alterations (GISTIC score < 0) were assigned as p53 mutant and CNV loss; The classified cell lines were further grouped based on the gene transcript levels: low (≤ 1st quartile), median (> 1st quartile and < 3rd quartile), high (≥ 3rd quartile). The effects of the mutation status or transcript levels on drug sensitivity were then determined with a linear model approach with log2 of the IC50 values as dependent variable and mutation status (Fig. 5B) or transcritpt levels (Fig. 5C-D and 5F-H) as independent variable.

### ChIP-Seq analysis

Reads from 30 p53 ChIP-seq datasets (**Supplementary Table** S10) were downloaded from the Sequence Read Archive (SRA). All datasets consisted of single ended Illumina reads. If multiple conditions were used in the same experiment, these were treated as separate datasets. Reads were trimmed using Trimmomatic version 0.32 (72) and bases with leading or trailing quality less than 3, across a 4 base sliding window with quality less than 15 were trimmed, as were Illumina adaptors. Reads with greater than 24 bases remaining were retained. Reads were mapped to hg38 using the BWA-mem alignment algorithm version 0.7.12 (73). The resulting BAM files were filtered to remove unmapped reads, duplicate reads (as identified with Picard MarkDuplicates 2.8.3 (http://broadinstitute.github.io/picard/) and reads with a mapping quality score less than 10. Peaks were called using MACS2 (version 2.1.1.20160309) (74) with the appropriate input dataset used as a control and a q-value cutoff of 0.01. This stringent threshold was selected to avoid overcalling peaks as a number of studies only had a single replicate for each condition. Insert size was estimated using the MACS2 predictd function. For datasets with multiple replicates, only peaks which were at least partially present in at least two replicates were maintained in the dataset.

### CRISPR/Cas9-mediated genome editing

The Cas9 expression vector was obtained from Addgene (#62988). sgRNAs were designed and constructed as described previously (75). Briefly, the sgRNA oligos were designed and analyzed using the CRISPR design tool (http://crispr.mit.edu/), and the ones with highest rating sores were selected. For the human U6 promoter-based transcription, a guanine (G) base was added to the 5′ of the sgRNA when the 20bp guide sequence did not begin with G. The oligo sequences for the sgRNA synthesis are listed in **Supplementary Table** S15. Next, the annealed oligos were cloned into the BbsI restriction sites of the Cas9 expression vector. The donor construct pMK-RQ-HDR-donor for generating the p53-REs knock in clones was synthesized by GeneArt Gene Synthesis service and integrated into the G418 resistant vector pMK-RQ (ThermoFisher). The donor construct rs78378222-HDR-donor was generated for the homology directed repair (HDR) in Hap1 cells. For genomic deletions, 5×10^5^ cells were seeded in a 12-well plate and transfected with 0.5 mg of each sgRNA constructs. After 24 hours, cells were incubated in puromycin for 48 hour. Subsequently, a single-cell suspension was prepared and seeded at a low density in 96-well plate for 3-4 weeks. Clones that were derived from more than one cell were excluded from further experiments. Individual colonies were picked and expanded for PCR-based genotyping with primers outside and inside of the targeting region (**Supplementary Table** S15). Correctly targeted clones were further confirmed by Sanger sequencing or TagMan genotyping, and the copy number of the heterozygous knock out cells was confirmed by TaqMan Copy Number Assays. For the knock-in, cells were transfected with a guide RNA (see sequences in **Supplementary Table** S15) together with a recombination donor flanked with 1-kb right and left homology arms where the PAM site was mutated to prevent donor DNA cleavage (a point mutation from CCA to GCA). Transfected cells were selected by treatment with puromycin and G418 for 48 hours. Based on the same procedures for genomic deletion, correctly targeted clones were validated by PCR-based genotyping, Sanger sequencing and copy number determination.

### Cell culture and their treatments

Testicular cancer cell lines TERA1, TERA2, 2102EP, Susa-CR, GH, were cultured in RPMI (Roswell Park Memorial Institute) medium containing 10% fetal bovine serum and 1% penicillin/streptomycin according to standard conditions. Susa cells were cultured in RPMI medium containing 20% fetal bovine and 1% penicillin/streptomycin. GCT27 and GCT27-CR were cultured in DMEM (Dulbecco’s modified Eagle’s medium) supplemented with 10% fetal bovine serum and 1% penicillin/streptomycin. Hap1 cells were obtained from Horizon Discovery Ltd and cultured in IMDM (Sigma-Aldrich Co Ltd) supplemented with 10% fetal bovine serum and 1% penicillin/streptomycin. FuGENE 6 Transfection Reagent (Promega) was used for DNA transfection. For transfection of siRNA, Lipofectamine RNAiMAX Transfection Reagent (ThermoFisher) was used.

### Drug screening

Cells were seeded in 384-well plates (flat bottom, black with clear bottom, Greiner) at density of about 2,000 cells per well in 81μl with cell dispenser (FlexDrop, PerkinElmer) and liquid handling robotics (JANUS, PerkinElmer) and incubated overnight. Next, library compounds (**Supplementary Table** S14) were added to a final concentration of 10μM, 1μM, 100nM or 10nM. Dasatinib (1uM) was added as positive control and DMSO (Vehicle, 0.1%) was added as negative control. After 72 hours, cell were fixed with 4% paraformaldehyde for 10 min, permeabilized with 0.5% Triton X-100 for 5 min, and then stained with 1:1000 dilution of 5mg/ml DAPI for 5 min. Next, the plates were imaged using a high-content analysis system (Operetta, PerkinElmer). The image data was analyzed by an image data storage and analysis system (Columbus, PerkinElmer). The cells with nuclear area>150 and nuclear intensity<700 were counted, and cell number was used as the viability readout.

### IC50 and combination index CI analyses

To determine an IC50, 8 multiply diluted concentrations (ranging from 0 to 10 μM for Nutlin3, 0 to 5 μM for Doxorubicin, 0 to 0.5 μM for Camptothecin and Topotecan, 0 to 20 μM for Dasatinib, Suntinib and Nilotinib, and 0 to 100 μM for Imatinib and Pazopanib) were used including a PBS control for 48 hour treatment and then cell viability was assessed by a MTT assay. The IC50 was calculated using the Graphpad Prism software. A constant ratio matrix approach was used to determine the combination index CI values (76). Single drug data and combination data was entered into Compusyn software (http://www.combosyn.com) to compute CI50 and dose-reduction index (DRI). CI50 is (CX/IC50(X)) + (CY/IC50(Y)), where (CX/IC50(X)) is the ratio of the drug X’s concentration (CX) in a 50% effective drug mixture to its 50% inhibitory concentration (IC50(X)) when applied alone. The CI50 values quantitatively depict synergistic (CI<1), additive (CI=1), and antagonistic effects (CI>1).

### *In vivo* study

All animal procedures were carried out under a Home Office licence (PPL30/3395), and mice were housed at Oxford University Biomedical Services, UK. 6-8 week-old female BALB/c nude mice (Charles River, UK) were injected subcutaneously with 5×10^6^ 2102EP cells in a 1:1 mixture of serum-free medium and Matrigel. When the average tumor volume reached approximately 130 mm^3^, animals were divided into four groups (6 per group) and received the following treatments: 1. Vehicle group: p.o. vehicle A (2% DMSO/ 30% PEG300/ dH2O) on day 1-5 & day 8-12, once daily; i.p. vehicle B (saline) on day 1, 8 and 12, once daily; 2. Doxorubicin group: i.p. Doxrubicin 4mg/ kg (Sigma) in vehicle B on day 1, 8 and 12, once daily; 3. Dasatinib group: p.o. Dasatinib 25mg/kg (Selleckchem) in vehicle A on day 1-5 and day 8-12, once daily; 4. Combination group: p.o. Dasatinib 25mg/kg on day 1-5 and day 8-12, once daily; i.p. Doxorubicin 4mg/ kg 1h after Dasatinib dosing on day 1, 8 and 12, once daily. Mouse weights and tumor volumes were measured 3 times per week. All mice were sacrificed on day 12, 2h after final treatments.

## Acknowledgments

This work was funded in part by the Ludwig Institute for Cancer Research, the Nuffield Department of Medicine, the Development Fund, Oxford Cancer Research Centre, University of Oxford, UK, by the Intramural Research Program of the National Institute of Environmental Health Sciences-National Institutes of Health (Z01-ES100475), and NIH grant (DP5-OD017937), US, and by the S-CORT Consortium from the Medical Research Council and Cancer Research UK.

## Declaration of Interests

The authors declare no competing interests.

## References

1. Dancey JE, Bedard PL, Onetto N, Hudson TJ. The genetic basis for cancer treatment decisions. Cell 2012;148(3): 409–20

2. Huang M, Shen A, Ding J, Geng M. Molecularly targeted cancer therapy: some lessons from the past decade. Trends Pharmacol Sci 2014;35(1): 41–50

3. Carter H, Marty R, Hofree M, Gross AM, Jensen J, Fisch KM, et al. Interaction Landscape of Inherited Polymorphisms with Somatic Events in Cancer. Cancer discovery 2017;7(4): 410–23

4. Lu C, Xie M, Wendl MC, Wang J, McLellan MD, Leiserson MD, et al. Patterns and functional implications of rare germline variants across 12 cancer types. Nature communications 2015;6:10086

5. Yurgelun MB, Chenevix-Trench G, Lippman SM. Translating Germline Cancer Risk into Precision Prevention. Cell 2017;168(4): 566–70

6. Stracquadanio G, Wang XT, Wallace MD, Grawenda AM, Zhang P, Hewitt J, et al. The importance of p53 pathway genetics in inherited and somatic cancer genomes. Nature Reviews Cancer 2016;16(4): 251–65

7. Martincorena I, Raine KM, Gerstung M, Dawson KJ, Haase K, Van Loo P, et al. Universal Patterns of Selection in Cancer and Somatic Tissues. Cell 2017;171(5): 1029–41 e21

8. Kastenhuber ER, Lowe SW. Putting p53 in Context. Cell 2017;170(6): 1062–78

9. Bouwman P, Jonkers J. The effects of deregulated DNA damage signalling on cancer chemotherapy response and resistance. Nature reviews Cancer 2012;12(9): 587–98

10. Lowe SW, Ruley HE, Jacks T, Housman DE. P53-Dependent Apoptosis Modulates the Cytotoxicity of Anticancer Agents. Cell 1993;74(6): 957–67

11. Weinstein JN, Myers TG, O’Connor PM, Friend SH, Fornace AJ Jr., Kohn KW, et al. An information-intensive approach to the molecular pharmacology of cancer. Science 1997;275(5298): 343–9

12. Kotler E, Shani O, Goldfeld G, Lotan-Pompan M, Tarcic O, Gershoni A, et al. A Systematic p53 Mutation Library Links Differential Functional Impact to Cancer Mutation Pattern and Evolutionary Conservation. Molecular cell 2018;71(1): 178–90 e8

13. Lang GA, Iwakuma T, Suh YA, Liu G, Rao VA, Parant JM, et al. Gain of function of a p53 hot spot mutation in a mouse model of Li-Fraumeni syndrome. Cell 2004;119(6): 861–72

14. Olive KP, Tuveson DA, Ruhe ZC, Yin B, Willis NA, Bronson RT, et al. Mutant p53 gain of function in two mouse models of Li-Fraumeni syndrome. Cell 2004;119(6): 847–60

15. Bykov VJN, Eriksson SE, Bianchi J, Wiman KG. Targeting mutant p53 for efficient cancer therapy. Nature reviews Cancer 2018;18(2): 89–102

16. Wade M, Li YC, Wahl GM. MDM2, MDMX and p53 in oncogenesis and cancer therapy. Nature reviews Cancer 2013;13(2): 83–96

17. Chen L, Agrawal S, Zhou W, Zhang R, Chen J. Synergistic activation of p53 by inhibition of MDM2 expression and DNA damage. Proceedings of the National Academy of Sciences of the United States of America 1998;95(1): 195–200

18. Hoe KK, Verma CS, Lane DP. Drugging the p53 pathway: understanding the route to clinical efficacy. Nature Reviews Drug Discovery 2014;13(3): 217–36

19. Suh YA, Post SM, Elizondo-Fraire AC, Maccio DR, Jackson JG, El-Naggar AK, et al. Multiple stress signals activate mutant p53 in vivo. Cancer Res 2011;71(23): 7168–75

20. Terzian T, Suh YA, Iwakuma T, Post SM, Neumann M, Lang GA, et al. The inherent instability of mutant p53 is alleviated by Mdm2 or p16INK4a loss. Genes & development 2008;22(10): 1337–44

21. Yue X, Zhao Y, Xu Y, Zheng M, Feng Z, Hu W. Mutant p53 in Cancer: Accumulation, Gain-of-Function, and Therapy. J Mol Biol 2017;429(11): 1595–606

22. Jansen R, Hottenga JJ, Nivard MG, Abdellaoui A, Laport B, de Geus EJ, et al. Conditional eQTL analysis reveals allelic heterogeneity of gene expression. Human molecular genetics 2017;26(8): 1444–51

23. Consortium GT. The Genotype-Tissue Expression (GTEx) project. Nature genetics 2013;45(6): 580–5

24. Gong J, Mei S, Liu C, Xiang Y, Ye Y, Zhang Z, et al. PancanQTL: systematic identification of cis-eQTLs and trans-eQTLs in 33 cancer types. Nucleic acids research 2018;46(D1):D971–D6

25. Cancer Genome Atlas N. Comprehensive molecular portraits of human breast tumours. Nature 2012;490(7418): 61–70

26. Cancer Genome Atlas Research N. Integrated genomic analyses of ovarian carcinoma. Nature 2011;474(7353): 609–15

27. Michailidou K, Lindstrom S, Dennis J, Beesley J, Hui S, Kar S, et al. Association analysis identifies 65 new breast cancer risk loci. Nature 2017;551(7678): 92–4

28. Phelan CM, Kuchenbaecker KB, Tyrer JP, Kar SP, Lawrenson K, Winham SJ, et al. Identification of 12 new susceptibility loci for different histotypes of epithelial ovarian cancer. Nature genetics 2017;49(5): 680–91

29. Melin BS, Barnholtz-Sloan JS, Wrensch MR, Johansen C, Il’yasova D, Kinnersley B, et al. Genome-wide association study of glioma subtypes identifies specific differences in genetic susceptibility to glioblastoma and non-glioblastoma tumors. Nature genetics 2017;49(5): 789–94

30. Stacey SN, Helgason H, Gudjonsson SA, Thorleifsson G, Zink F, Sigurdsson A, et al. New basal cell carcinoma susceptibility loci. Nature communications 2015;6:6825

31. Basu S, Murphy ME. Genetic Modifiers of the p53 Pathway. Cold Spring Harb Perspect Med 2016;6(4):a026302

32. Milne RL, Kuchenbaecker KB, Michailidou K, Beesley J, Kar S, Lindstrom S, et al. Identification of ten variants associated with risk of estrogen-receptor-negative breast cancer. Nature genetics 2017;49(12): 1767–78

33. Stacey SN, Sulem P, Jonasdottir A, Masson G, Gudmundsson J, Gudbjartsson DF, et al. A germline variant in the TP53 polyadenylation signal confers cancer susceptibility. Nature genetics 2011;43(11): 1098–103

34. Burckstummer T, Banning C, Hainzl P, Schobesberger R, Kerzendorfer C, Pauler FM, et al. A reversible gene trap collection empowers haploid genetics in human cells. Nature methods 2013;10(10): 965–71

35. Liberzon A, Birger C, Thorvaldsdottir H, Ghandi M, Mesirov JP, Tamayo P. The Molecular Signatures Database (MSigDB) hallmark gene set collection. Cell Syst 2015;1(6): 417–25

36. Gurtner A, Starace G, Norelli G, Piaggio G, Sacchi A, Bossi G. Mutant p53-induced up-regulation of mitogen-activated protein kinase kinase 3 contributes to gain of function. The Journal of biological chemistry 2010;285(19): 14160–9

37. Solomon H, Dinowitz N, Pateras IS, Cooks T, Shetzer Y, Molchadsky A, et al. Mutant p53 gain of function underlies high expression levels of colorectal cancer stem cells markers. Oncogene 2018;37(12): 1669–84

38. Robles AI, Harris CC. Clinical outcomes and correlates of TP53 mutations and cancer. Cold Spring Harbor perspectives in biology 2010;2(3):a001016

39. Hientz K, Mohr A, Bhakta-Guha D, Efferth T. The role of p53 in cancer drug resistance and targeted chemotherapy. Oncotarget 2017;8(5): 8921–46

40. Fischer M. Census and evaluation of p53 target genes. Oncogene 2017;36(28): 3943–56

41. Iorio F, Knijnenburg TA, Vis DJ, Bignell GR, Menden MP, Schubert M, et al. A Landscape of Pharmacogenomic Interactions in Cancer. Cell 2016;166(3): 740–54

42. Lennartsson J, Ronnstrand L. Stem cell factor receptor/c-Kit: from basic science to clinical implications. Physiological reviews 2012;92(4): 1619–49

43. Zeron-Medina J, Wang X, Repapi E, Campbell MR, Su D, Castro-Giner F, et al. A polymorphic p53 response element in KIT ligand influences cancer risk and has undergone natural selection. Cell 2013;155(2): 410–22

44. Turnbull C, Rapley EA, Seal S, Pernet D, Renwick A, Hughes D, et al. Variants near DMRT1, TERT and ATF7IP are associated with testicular germ cell cancer. Nature genetics 2010;42(7):604–U178

45. Litchfield K, Levy M, Orlando G, Loveday C, Law PJ, Migliorini G, et al. Identification of 19 new risk loci and potential regulatory mechanisms influencing susceptibility to testicular germ cell tumor. Nature genetics 2017;49(7):1133–+

46. Vogelstein B, Lane D, Levine AJ. Surfing the p53 network. Nature 2000;408(6810): 307–10

47. Pommier Y. Topoisomerase I inhibitors: camptothecins and beyond. Nature reviews Cancer 2006;6(10): 789–802

48. Flaherty KT, Hodi FS, Fisher DE. From genes to drugs: targeted strategies for melanoma. Nature reviews Cancer 2012;12(5): 349–61

49. Litchfield K, Levy M, Huddart RA, Shipley J, Turnbull C. The genomic landscape of testicular germ cell tumours: from susceptibility to treatment. Nat Rev Urol 2016;13(7): 409–19

50. Noel EE, Yeste-Velasco M, Mao X, Perry J, Kudahetti SC, Li NF, et al. The association of CCND1 overexpression and cisplatin resistance in testicular germ cell tumors and other cancers. The American journal of pathology 2010;176(6): 2607–15

51. Koster R, di Pietro A, Timmer-Bosscha H, Gibcus JH, van den Berg A, Suurmeijer AJ, et al. Cytoplasmic p21 expression levels determine cisplatin resistance in human testicular cancer. The Journal of clinical investigation 2010;120(10): 3594–605

52. Vazquez A, Bond EE, Levine AJ, Bond GL. The genetics of the p53 pathway, apoptosis and cancer therapy. Nat Rev Drug Discov 2008;7(12): 979–87

53. Kruiswijk F, Labuschagne CF, Vousden KH. p53 in survival, death and metabolic health: a lifeguard with a licence to kill. Nat Rev Mol Cell Bio 2015;16(7): 393–405

54. Kerns SL, Fung C, Monahan PO, Ardeshir-Rouhani-Fard S, Abu Zaid MI, Williams AM, et al. Cumulative Burden of Morbidity Among Testicular Cancer Survivors After Standard Cisplatin-Based Chemotherapy: A Multi-Institutional Study. J Clin Oncol 2018:JCO2017770735

55. Ravandi F, O’Brien S, Thomas D, Faderl S, Jones D, Garris R, et al. First report of phase 2 study of dasatinib with hyper-CVAD for the frontline treatment of patients with Philadelphia chromosome-positive (Ph+) acute lymphoblastic leukemia. Blood 2010;116(12): 2070–7

56. Grobner SN, Worst BC, Weischenfeldt J, Buchhalter I, Kleinheinz K, Rudneva VA, et al. The landscape of genomic alterations across childhood cancers. Nature 2018;555(7696): 321–7

57. PCAWG Germline Working group. Germline determinants of the somatic mutation landscape in 2,642 cancer genomes. bioRxiv 2017

58. Saiki AY, Caenepeel S, Cosgrove E, Su C, Boedigheimer M, Oliner JD. Identifying the determinants of response to MDM2 inhibition. Oncotarget 2015;6(10): 7701–12

59. Alt JR, Greiner TC, Cleveland JL, Eischen CM. Mdm2 haplo-insufficiency profoundly inhibits Myc-induced lymphomagenesis. The EMBO journal 2003;22(6): 1442–50

60. Hohenstein P. Tumour suppressor genes--one hit can be enough. PLoS biology 2004;2(2):E40

61. Muller PA, Vousden KH. Mutant p53 in cancer: new functions and therapeutic opportunities. Cancer cell 2014;25(3): 304–17

62. Bailey MH, Tokheim C, Porta-Pardo E, Sengupta S, Bertrand D, Weerasinghe A, et al. Comprehensive Characterization of Cancer Driver Genes and Mutations. Cell 2018;174(4): 1034–5

63. Chang MT, Asthana S, Gao SP, Lee BH, Chapman JS, Kandoth C, et al. Identifying recurrent mutations in cancer reveals widespread lineage diversity and mutational specificity. Nature biotechnology 2016;34(2): 155–63

64. Bouaoun L, Sonkin D, Ardin M, Hollstein M, Byrnes G, Zavadil J, et al. TP53 Variations in Human Cancers: New Lessons from the IARC TP53 Database and Genomics Data. Human mutation 2016;37(9): 865–76

65. Giacomelli AO, Yang X, Lintner RE, McFarland JM, Duby M, Kim J, et al. Mutational processes shape the landscape of TP53 mutations in human cancer. Nature genetics 2018;50(10): 1381–7

66. Willer CJ, Li Y, Abecasis GR. METAL: fast and efficient meta-analysis of genomewide association scans. Bioinformatics 2010;26(17): 2190–1

67. Chang CC, Chow CC, Tellier LC, Vattikuti S, Purcell SM, Lee JJ. Second-generation PLINK: rising to the challenge of larger and richer datasets. Gigascience 2015;4:7

68. Cancer Genome Atlas Research N, Weinstein JN, Collisson EA, Mills GB, Shaw KR, Ozenberger BA, et al. The Cancer Genome Atlas Pan-Cancer analysis project. Nature genetics 2013;45(10): 1113–20

69. Korn JM, Kuruvilla FG, McCarroll SA, Wysoker A, Nemesh J, Cawley S, et al. Integrated genotype calling and association analysis of SNPs, common copy number polymorphisms and rare CNVs. Nature genetics 2008;40(10): 1253–60

70. Das S, Forer L, Schonherr S, Sidore C, Locke AE, Kwong A, et al. Next-generation genotype imputation service and methods. Nature genetics 2016;48(10): 1284–7

71. Liu JF, Lichtenberg T, Hoadley KA, Poisson LM, Lazar AJ, Cherniack AD, et al. An Integrated TCGA Pan-Cancer Clinical Data Resource to Drive High-Quality Survival Outcome Analytics. Cell 2018;173(2):400–+

72. Bolger AM, Lohse M, Usadel B. Trimmomatic: a flexible trimmer for Illumina sequence data. Bioinformatics 2014;30(15): 2114–20

73. Li H, Durbin R. Fast and accurate short read alignment with Burrows– Wheeler transform. Bioinformatics 2009;25(14): 1754–60

74. Zhang Y, Liu T, Meyer CA, Eeckhoute J, Johnson DS, Bernstein BE, et al. Model-based Analysis of ChIP-Seq (MACS). Genome Biology 2008;9(9):R137

75. Ran FA, Hsu PD, Wright J, Agarwala V, Scott DA, Zhang F. Genome engineering using the CRISPR-Cas9 system. Nature protocols 2013;8(11): 2281–308

76. Chou TC. Theoretical basis, experimental design, and computerized simulation of synergism and antagonism in drug combination studies. Pharmacol Rev 2006;58(3): 621–81

