## Supplementary Methods and Figures S1-S7 for "Germline and somatic genetic variants in the p53 pathway interact to affect cancer risk, progression and drug response"

#### Supplementary Materials

Supplementary Methods

Legends for Supplemental Figures S1-S7

Supplemental Figures S1-S7

#### Supplementary Methods

##### SDS-PAGE and western blotting

SDS-PAGE and western blotting was performed as described in (1). . The antibodies against p53 (DO-1), c-KIT (E-1), PARP1 (H-250), and β-Actin (C4) were from Santa Cruz (Dallas, TX, USA). The antibodies against acetylated p53 (K382), cleaved Caspase 3 (Asp175) were from Cell Signalling (Danvers, MA, USA). HRP-coupled secondary antibodies were from Dako (Darmstadt, Germany).

##### RNA isolation, cDNA synthesis and Quantitative real-time reverse transcription PCR

Cells were lysed and total RNA was prepared using the RNeasy kit (Qiagen) according to the manufacturer's protocol. cDNA was synthesized using SuperScript VILO Master Mix (ThermoFisher). Quantitative real-time reverse transcription PCR (qRT-PCR) was carried out in triplicates with SYBR® Green Real-Time PCR Master Mix (Qiagen) on an ABI StepOnePlus System (Applied Biosciences). The transcript levels were normalised by the readings for GAPDH.

##### MTT assay

Cells were plated into a 96-well plate overnight. MTT (3-(4,5-Dimethylthiazol-2-yl)-2,5-Diphenyltetrazolium Bromide) (ThermoFisher) was dissolved in PBS and added to the cells to a final concentration of 0.75 mg/ml for 4h. Thereafter, the medium was aspirated. The formazan salt was solubilised in isopropanol and measured at 570 nm using an ELISA reader.

##### Cell proliferation and migration

The proliferation and migration rates of the cells were determined by using the xCELLigence system (2). Specifically, for cell proliferation, 3,000 cells were seeded in the 16-well E-Plate incorporated with sensor electrode arrays, and the cell index, which is an impedance-based readout of cell numbers, was continuously monitored for 150 hours. For cell migration, 30,000 cells were seeded in the 16-well CIM-Plate with a lower chamber and upper chamber. The upper chamber contains a microporous membrane (8 µm pore diameter) with electrode arrays on the bottom side. The real-time cell index readouts were monitored for 35 hours.

**Genetic copy number determination**

Copy numbers of the p53-bound *KITLG* region were determined using custom TaqMan Copy Number Assay (assay ID: chr12SCF_CDFVKNN; target location: hg38 chr12:88559379-88559679; FAM labeled MGB probe) and normalized to a TaqMan Copy Number Reference Assay RNase P with a VIC-labeled TAMRA probe (ThermoFisher). Briefly, a TaqMan Copy Number Assay, a TaqMan Copy Number Reference Assay, a Genotyping Master Mix, and a gDNA sample were mixed together in a single well of a 96-well plate. Following the manufacturer's instructions, all qPCR reactions were performed in triplicate on a StepOnePlus Real-Time PCR System (ThermoFisher).

**References**

1. Zhang P, Elabd S, Hammer S, Solozobova V, Yan H, Bartel F, et al. TRIM25 has a dual function in the p53/Mdm2 circuit. Oncogene 2015;34(46):5729-38
2. Ke N, Wang X, Xu X, Abassi YA. The xCELLigence system for real-time and label-free monitoring of cell viability. Methods in molecular biology 2011;740:33-43

#### Supplementary figure legends

**Figure S1.** (A) A scatter plot of the fold enrichment of eGenes on the x-axis (log2 scale), and the adjusted p-value on the y-axis (-log10 scale), amongst each pathways relative to all eGenes of the genome. The enrichment of eGenes in p53 pathway genes is marked in red and the other 185 annotated KEGG pathways are marked in grey. The horizontal dashed lines represent the FDR-adjusted p value of 0.05. (B) Allelic discrimination plots of the p53 poly(A) SNP using TaqMan genotyping assays on CRISPR/Cas9-edited Hap1 clones. Grey dots represent genotype with A allele, and red and orange dots represent genotype with C allele. (C) Kaplan-Meier survival curves for PFI or OS in patients (left panel: 6,977 pan-cancer patients for PFI; middle panel: 6,979 pan-cancer patients for OS; right panel: 712 breast cancer patients for PFI;) carrying either the major or the minor allele of the p53 poly(A) SNP. Below each plot, the number of patients for each time point, and genotype class, are indicated.

**Figure S2.** (A) Genetic fine mapping with an independent TGCT GWAS cohorts identified six p53-bound SNPs with the strongest TGCT GWAS signal (high $-\text{Log}_{10}$ p-values) and which are in high linkage disequilibrium in Europeans ($r^2 > 0.95$; highlighted in red square). (B) A bar plot showing the number of the patients with high-stage (IS, II, III) or low low-stage (I) TGCT based on American Joint Committee on Cancer (AJCC) pathology stages if available, and the clinical stages for those without pathological stage (also see **Supplementary Table** S12).

**Figure S3. Characterization of the p53-bound *KITLG* locus in non-edited p53-REs+/+, knock-out p53-REs-/- or knock-in clones.** (A) A diagram of the CRISPR/cas9-mediated genomic editing utilized. Red and green arrows indicate the primers for PCR-based validation. (B) PCR results of genomic DNA isolated from p53-REs+/+ or p53-REs-/- clones by using primers indicated in (A) where two sets of primers (see **Supplementary Table** S15 for primer sequences) were designed for genomic editing detection of the sgRNAs-targeted *KITLG* locus in TERA1 and TERA2 cells. (C) Sanger sequencing analysis of the non-edited p53-REs+/+ and knock-out p53-REs-/- clones. The 5′-NGG adjacent motifs (PAMs) are shown in green and sites of cleavage by Cas9 are indicated by red triangles. (D) PCR-based validation of non-edited p53-REs+/+ or heterozygous knock-out p53-REs+/- clones in Susa-CR and GH cells. (E) Copy numbers of the p53-bound *KITLG* region were measured using custom TaqMan Copy Number Assays. Error bars represent SEM of at least 2 independent biological replicates. (F) Representative sequencing chromatograms for p53-REs+/- clones with non-risk haplotype (upper panel) or risk haplotype (bottom panel). Each pair of alleles is highlighted in purple bars. (G) A diagram of the CRISPR-mediated genomic editing utilized for generating the knock-in clones. The 5′-NGG adjacent motifs (PAMs) are shown in orange. The primers used for PCR based genotyping and sanger sequencing are shown by arrows (see Table S11 for primer sequences). (H) PCR results of genomic DNA isolated from knock out or knock in clones by using primers indicated in (G) where three sets of primers were designed for genomic editing detection of the sgRNAs-targeted *KITLG* locus. (I). Sanger sequencing analysis of the non-edited and the knock-in clones. In the knock-in clones, the p53-bound region in *KITLG* intron 1 was fully inserted by using a DNA donor containing a mutated PAM site (a point mutation from CCA to GCA).

**Figure S4. The p53-bound cancer risk region is a p53-regulated *KITLG* enhancer.** (A) Differences in cDNA levels of transcripts encoded by genes surrounding the *KITLG* p53-RE cluster (±2Mbp). cDNA levels were measured using qRT-PCR normalized to GAPDH and error bars represent SEM of three independent experiments. p-values were calculated using a two-tailed t-test. (B-C) Dot plots of *KITLG* cDNA levels that were measured using qRT-PCR and normalized to GAPDH as described above. In total, 2 to 3 clones of each genotype were analyzed in 3 independent biological replicates (Susa-CR cells) and 2 independent biological replicates (GH cells).

**Figure S5. Knock down of c-KIT by siRNAs or deletion of the p53 bound region of *KITLG* reduces cell proliferation and migration.** (A-B) Growth curves of TERA1 and TERA2 cells that were transfected with two different siRNAs targeted against c-KIT or with a control non-targeted siRNA pool. Real-time monitoring of

TERA1 and TERA2 cell proliferation or migration by xCELLigence RTCA Systems (see STAR Methods). Cells were treated with either a non-target control siRNA pool or two different siRNAs targeting c-KIT (c-KIT siRNA#1 and c-KIT siRNA#2), and seeded in the E-Plate for the determination of cell proliferation. 48 h after transfection, cells were seeded in CIM-Plate for the determination of cell migration. Cell index readings were continuously monitored. Error bars represent SEM of at least 2 independent experiments. p-values were calculated by two-way ANOVA followed by Tukey's multiple comparison test. (C-D) Growth curves of p53-REs+/+ clones (grey bars), p53-REs-/- clones (red bars) of TERA1 and TERA2 cells. At least two clones of p53-REs+/+ and p53-REs-/- TERA1 and TERA2 cells were seeded and determined for cell proliferation and migration. Error bars represent SEM of at least 2 independent experiments. p-values were calculated by two-way ANOVA followed by Tukey's multiple comparison test.

**Figure S6. The p53 bound region of *KITLG* protects against p53-mediated apoptosis.** (A) A bar graphs of average IC50 values of TERA1 and TERA2 cells that were transfected with two different siRNAs targeted against c-KIT or with a control non-targeted siRNA pool. The cell viability was analyzed by MTT assay, and IC50 calculated. Error bars represent SEM in 4 independent biological replicates. p-values were calculated by the two-tailed t-test. (B) Western blot analysis of the siRNA treated cells with or without Nutlin3 treatment for 6 hours. The cells were lysed and analyzed for c-KIT, p53 and cleaved-caspase3 protein expression. (C) Western blot analysis of TERA1 or TERA2 cells that were treated with or without Nutlin3 for 6 hours, lysed and analyzed for p53, acetylated p53, Parp1 and cleaved-caspase3 protein expression. (D) Western blot analysis of p53-REs+/+, p53-REs-/- and -/KI of TERA1 cells that were treated with an increasing dose of nutlin3 for 24 hours. The cells were lysed and analyzed for p53, Parp1 and cleaved-caspase3 protein expression. (E) Western blot analysis of the siRNA treated cells with or without Nutlin3 treatment for 6 hours. The cells were lysed and analyzed for p53 and cleaved-caspase3 protein expression. (F) Dose-response curves for cell viability of the siRNA treated cells with or without Nutlin3 treatment. Error bars represent SEM of 2 different clones per genotype.

**Figure S7. p53/*KITLG* pro-survival signaling can attenuate responses to p53-activating agents.** (A) Schematic overview for the microscopy-based high-content drug screening. (B) Scatter plots of the ratio (log10 scale) of cell viability between p53-REs+/+ and p53-REs-/- cells in response to different compounds. The compounds to which the p53-REs -/- cells respond better (blue dots) are defined as hits (i.e., the relative cell viability after treatment in p53-REs+/+ cells versus p53-REs-/- cells is more than1.5 fold in both replicates). Venn diagram (right panel) of the overlap between the number of hits in TERA1 and TERA2 cells. p-value for the statistical significance of the overlap was calculated by hypergeometric probability test (http://nemates.org/MA/progs/overlap_stats.html). (C) Bar plots of the IC50 values of TERA1 and TERA2 cells in response to 5 different c-KIT inhibitors. Error bars represent SD of 3 independent biological replicates. (D-E) Bar plots of combination indexes of Topotecan, Doxorubicin, Cisplatin or Camptothecin with Dasatinib in p53-REs+/+ (grey bars) and p53-REs-/- (blue bars) of TERA1 and TERA2 cells. (F) Weight curves of non-tumor bearing female BALB/c mice upon combination treatment of Doxorubicin and Dasatinib. Mice body were weighed daily for the duration of the study. Error bars represent means ± SEM (n=3). p.o. Dasatinib (25 mg/kg) on day 1-5; i.p. Dxorubicin (4 mg/kg) 1h after Dasatinib dosing on day 1. p.o.: oral administration; i.p.: intraperitoneal injection. On day 15, all mice were sacrificed.

### Supplementary figure 1

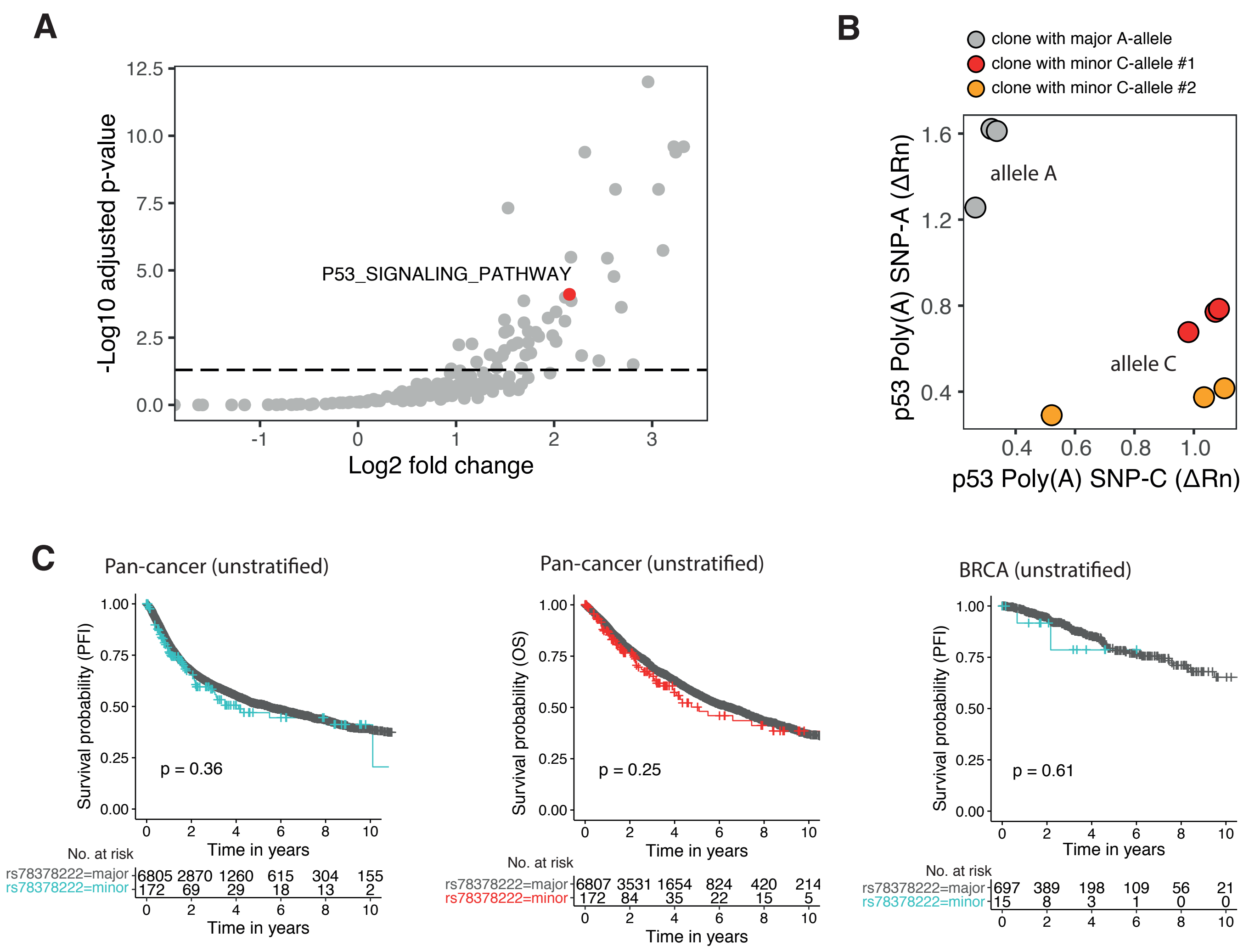

### Supplementary figure 2

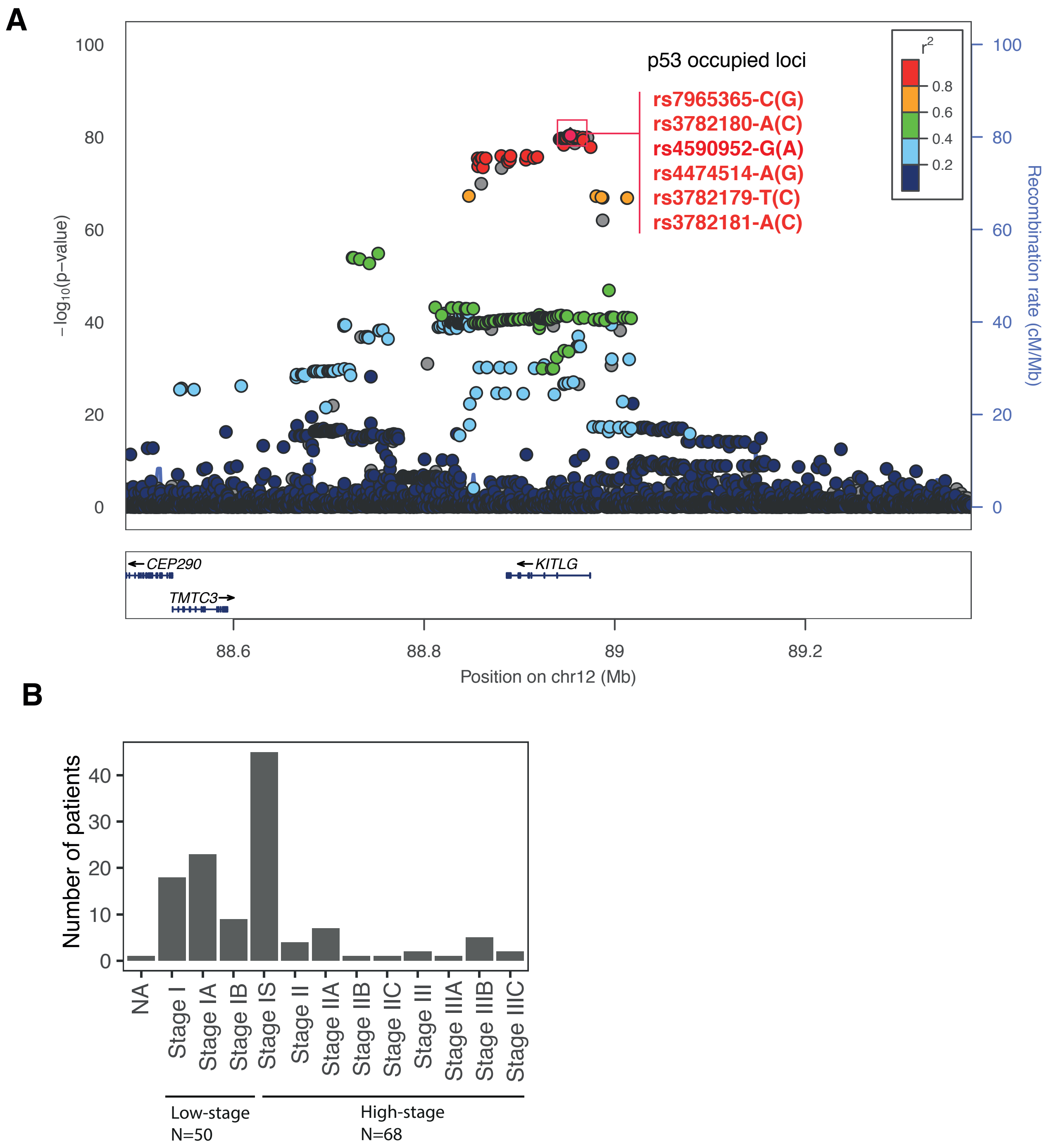

### Supplementary figure 3

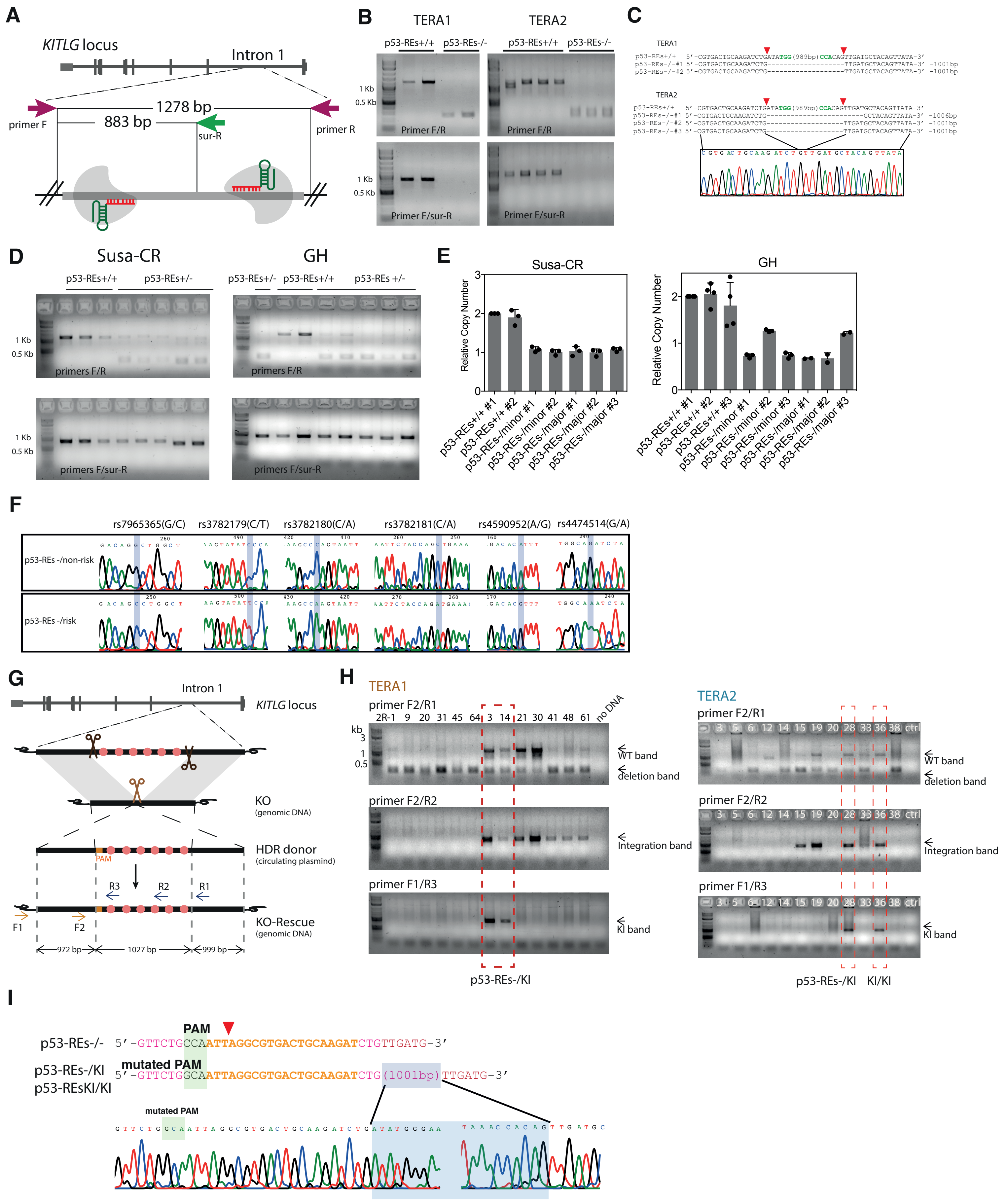

### Supplementary figure 4

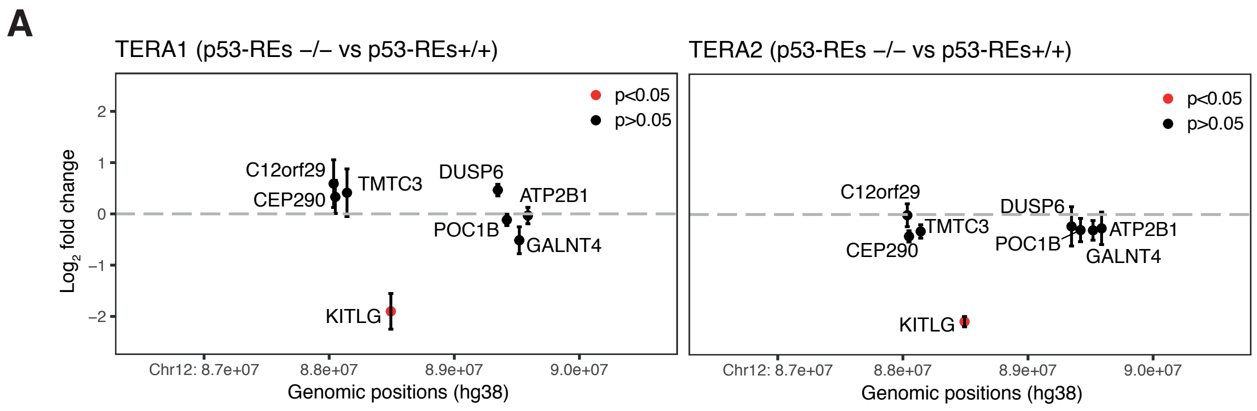


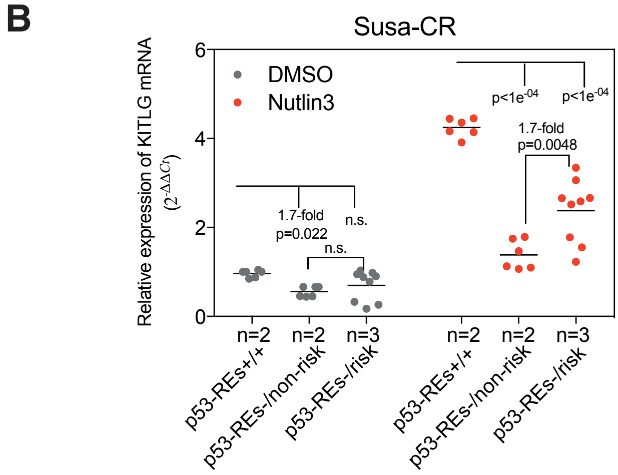


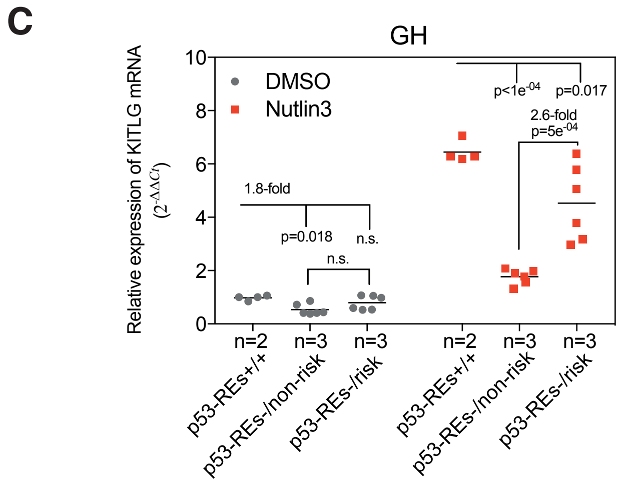

### Supplementary figure 5

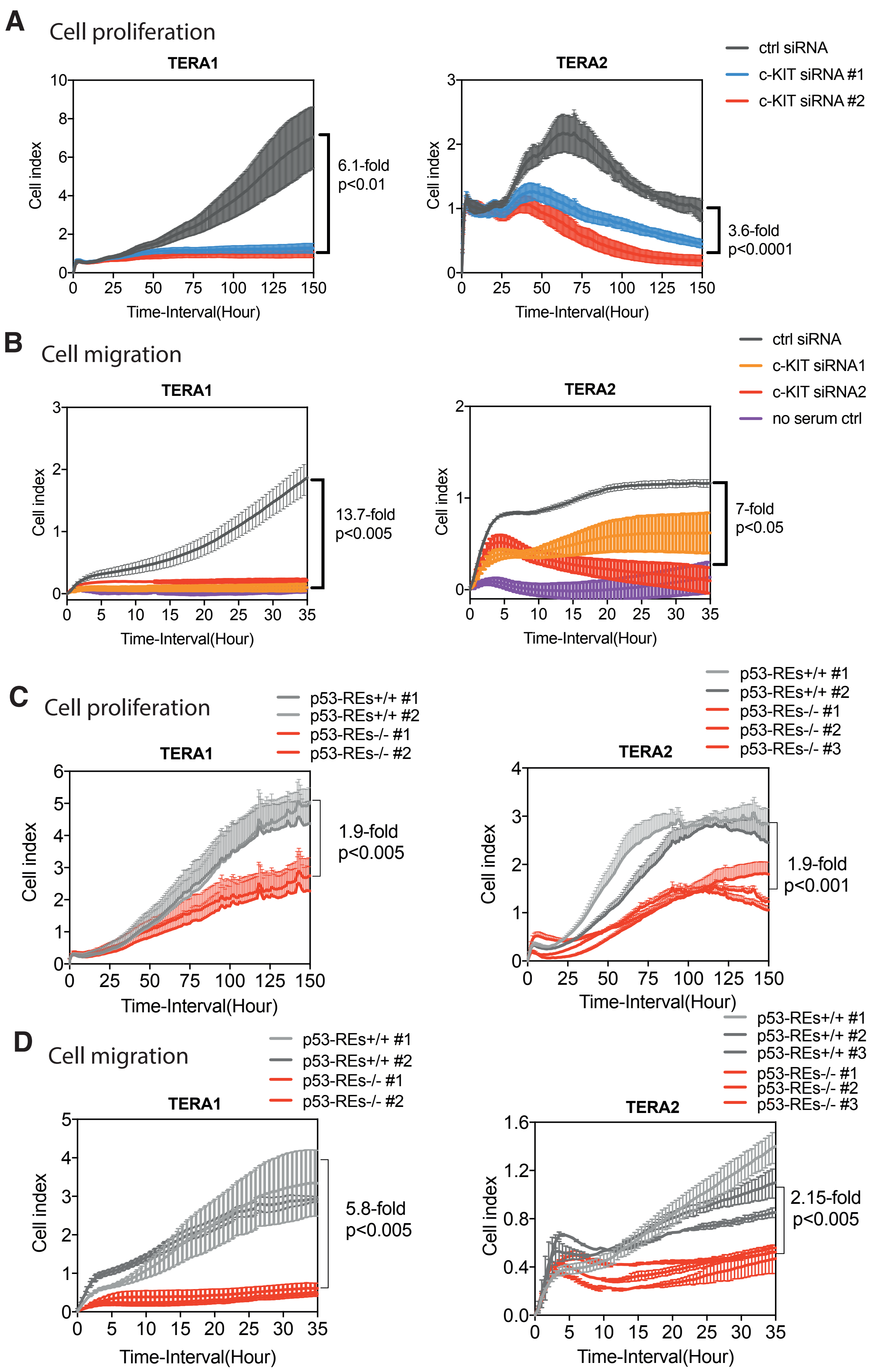

### Supplementary figure 6

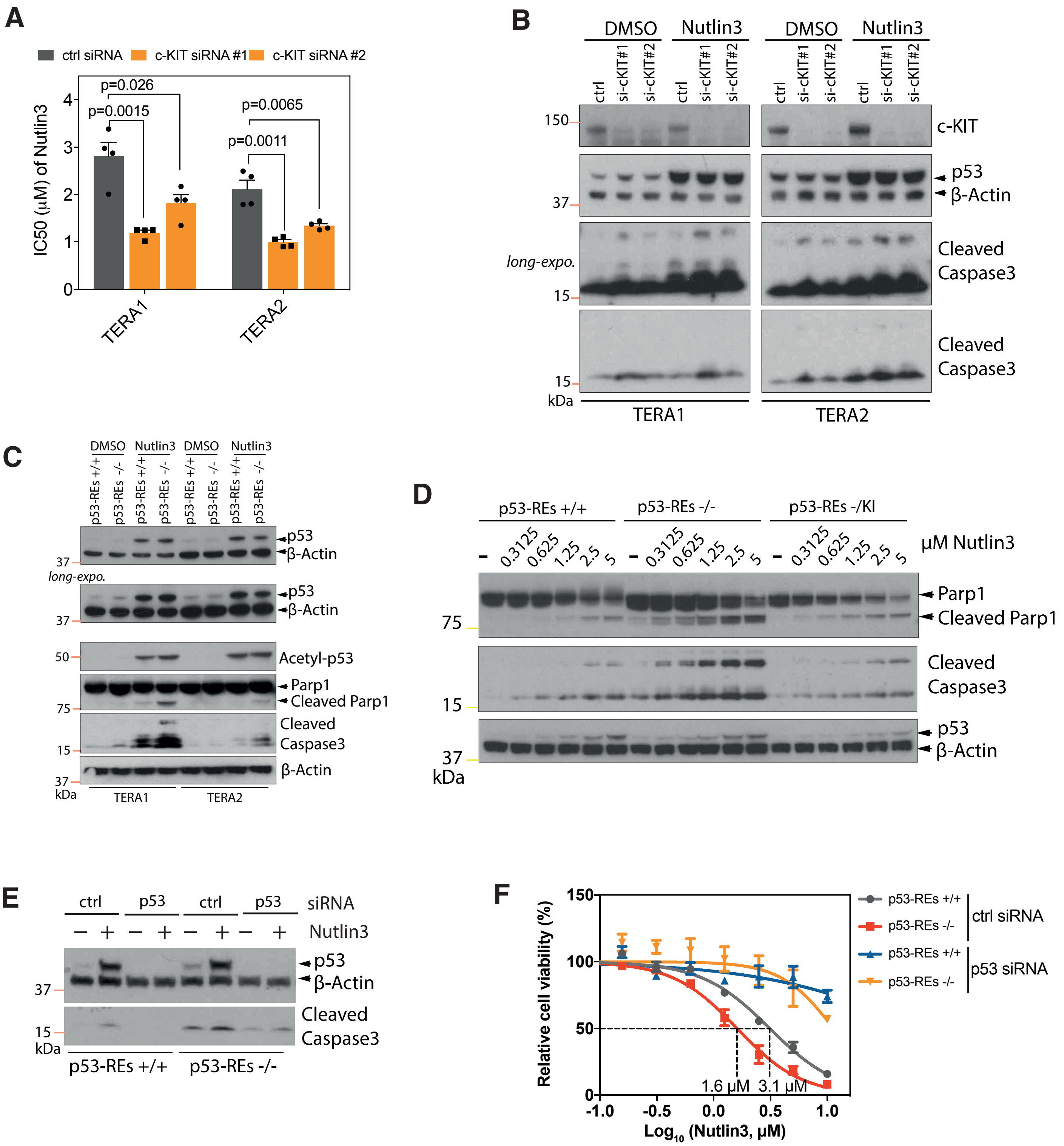

### Supplementary figure 7

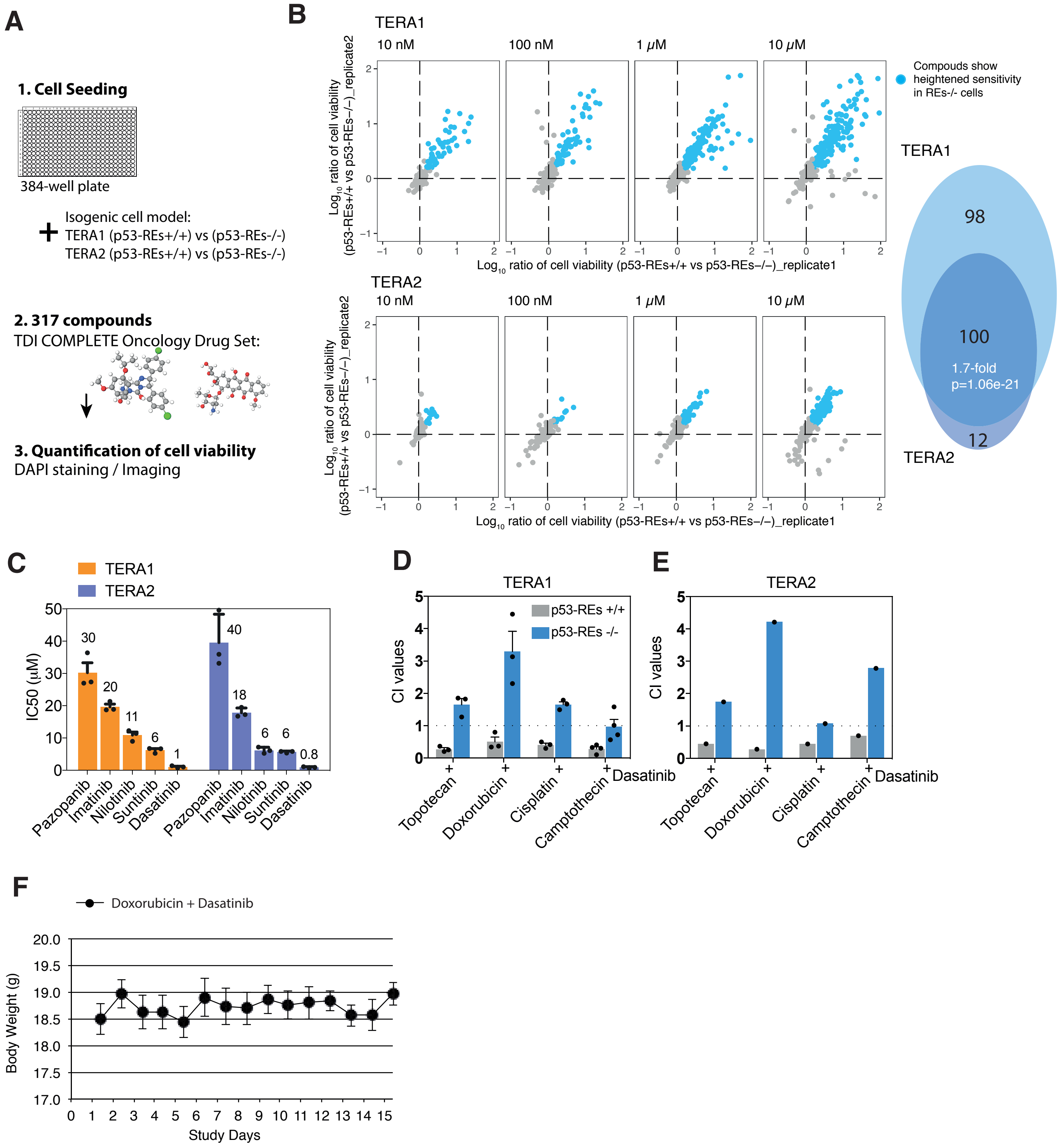
